# *Plasmodium falciparum* Raf kinase inhibitor is a lipid binding protein that interacts with CDPK1 and regulates its activity in asexual blood stage

**DOI:** 10.1101/2023.08.23.554426

**Authors:** Manish Sharma, Deepak Krishnan, Pooja Negi, Komal Rani, Amjesh Revikumar, Manoj Munde, Abhisheka Bansal

**Author notes:** Center for Systems Biology and Molecular Medicine (CSBMM), Yenepoya Research Centre (YRC), Yenepoya (Deemed to be University), Mangalore 575018, India. Institute of Biological Chemistry, Academia Sinica, Taiwan, National Taiwan University, Taipei, Taiwan (Komal Rani).

## Abstract

Raf Kinase Inhibitor Protein (RKIP) is an important regulator of MAPK signaling pathway in multicellular eukaryotes. *Plasmodium falciparum* RKIP (PfRKIP) is a putative phosphatidylethanolamine binding protein (PEBP) that shares limited similarity with *Homo sapiens* RKIP (HsRKIP). Interestingly, critical components of MAPK pathway are not expressed in malaria parasite and the physiological function of PfRKIP remains unknown. PfRKIP is expressed throughout the asexual schizogony with maximum expression in late schizonts. Interestingly, PfRKIP and HsRKIP show pH dependent differential interaction profiles with various lipids. At physiological pH, PfRKIP show interaction with PE and lipids containing phosphorylated phosphatidylinositol group; however, HsRKIP show no interaction under the same conditions. Mutation of conserved residues in the PEBP domain of PfRKIP decreases its interaction with PI(3)P. Furthermore, our results suggest that PfRKIP leads to increase in the autophosphorylation of PfCDPK1 that leads to transphosphorylation of substrates by PfCDPK1. Using various *in vitro* and *in vivo* experiments we have demonstrated the interaction of PfRKIP with PfCDPK1 and have also identified key residues in PfRKIP that play important role in this interaction. Interestingly, locostatin, a specific inhibitor of mammalian RKIP increased the interaction of PfRKIP with PfCDPK1 that perhaps leads to the sequestration of PfCDPK1 in a heterodimeric complex. Importantly, treatment of malaria parasite with locostatin shows dose dependent inhibition of parasite growth. This study suggests that specific inhibitors that modify PfRKIP leading to increase in its interaction with PfCDPK1 may be designed and explored as novel anti-malarial compounds to inhibit malaria parasite growth.

## INTRODUCTION

Malaria is a major infectious disease of developing nations. Five species of malaria parasite, *Plasmodium* are known to infect humans out of which, *P*. *falciparum* and *P*. *vivax* are majorly responsible for malaria related mortality and morbidity. Effective treatment is available against clinical infections of malaria however, there is growing concern of emerging resistance by malaria parasite against front-line drugs. Therefore, heightened global efforts are required for discovery of new anti-malarial drugs. Understanding of the molecular basis of key biological processes in the parasite lifecycle is pivotal for design and development of novel compounds with anti-malarial activity.

A phosphatidylethanolamine binding protein (PEBP) was initially isolated from bovine brain and was shown to bind phosphatidylethanolamine using radioactive PE [1, 2]. PEBP domain containing proteins are present across all prokaryotic and eukaryotic organisms. PEBP family members show tissue specific expression pattern in mammals [3] and are known to mediate critical processes including membrane fluidity and biogenesis [4–6] secretion of acetylcholine during neuronal development [6, 7], inhibition of serine proteases [8], and spermiogenesis/sperm maturation [9]. The PEBP domain containing family in *P*. *falciparum* is constituted by two members: PF3D7_0303900 and PF3D7_1219700. PF3D7_1219700 encodes a raf kinase inhibitor protein (RKIP) that is known to regulate the MAPK and NFκB signalling pathways in multicellular eukaryotes [10, 11] and hence play an important role in control of cellular proliferation, differentiation, and transformation: critical processes that are directly involved in regulation of tumorigenesis [12]. PfRKIP was reported as a putative phosphatidylethanolamine binding protein way back in 1995. Although, two putative homologs of MAPKs have been reported in *P*. *falciparum* however the canonical MAP3K and MEK are not represented [13–15] leading to a suggestion that the classical 3 component Raf1/MEK/ERK signalling cascade is perhaps not present in the malaria. Under this situation, the physiological function of RKIP in the malaria parasite biology remains elusive. PfRKIP was shown to downregulate the transphosphorylation activity of PfCDPK1 through *in vitro* kinase assays. However, the mechanistic understanding of regulation of kinase activity of PfCDPK1 by PfRKIP remains unexplored. Furthermore, we are not aware of any study that has demonstrated the phospholipid binding ability of PfRKIP. Additionally, the amino acids in PfRKIP that are important molecular determinants for the lipid and substrate binding activity of PfRKIP have not been studied.

Here, we have performed the biochemical characterization of PfRKIP and shown that the recombinant PfRKIP interacts with different lipids in a pH dependent manner. We have shown that PfRKIP binds phosphatidylethanolamine at the physiological pH along with other physiologically relevant lipids belonging to phosphorylated phosphatidylinositols. We have also determined the key residues in PfRKIP that play important role in interaction of PfRKIP with lipids. Importantly, here we have demonstrated that PfCDPK1 interacts with PfRKIP and established their co-localization within segmented schizonts and merozoites through co-immunoprecipitation and immunofluorescence assay, respectively. Furthermore, our results suggest that PfRKIP could play a crucial role in regulating the kinase activity of PfCDPK1. Finally, we have shown that treatment of late stage parasites with locostatin, a known pharmacological inhibitor of mammalian RKIP kills the parasite by blocking the invasion of red blood cells.

## RESULTS

### PfRKIP is expressed throughout the asexual blood stages

We tested the expression of *pfrkip* in the asexual blood stages of *P*. *falciparum* using semi-quantitative reverse transcriptase PCR, Western blot and immunofluorescence microscopy. RNA was extracted from highly synchronized ring (9-13 HPI), trophozoite (32-36 HPI), and schizont (44-48 HPI) stages and used for preparing complementary DNA (cDNA). The cDNA from each stage was used to amplify *pfrkip* with gene specific primers, RKIPFRT and RKIPRRT. Amplicon of desired size, 118 bp was obtained in all the stages (Fig. 1a). The expression of *pfrkip* in each stage was normalized with *pfgapdh*, amplified with GAPDHRTF/ GAPDHRTR, for the same stage and plotted as a histogram from two independent biological experiments (Fig. 1a). Maximum amplification was observed in schizonts followed by rings and trophozoites (Fig. 1a). We further tested the expression of PfRKIP protein through Western blot. Lysates from highly synchronized ring, trophozoite and schizont stage parasites were prepared and probed with in-house generated anti-PfRKIP antibody. A band corresponding to the size of full length PfRKIP (~ 21.5 kDa) was obtained in all the stages with maximum intensity in the schizonts followed by trophozoites and rings (Fig. 1b). Asynchronous parasite lysate probed with pre-immune sera did not result in any signal on the Western blot at the position corresponding to PfRKIP (Fig. S1b). Moreover, the pre-immune sera did not recognize recombinant PfRKIP that was otherwise readily recognized by the anti-PfRKIP antibodies (Fig. S1a). To further verify the expression of PfRKIP in blood stages, we generated a transgenic parasite where the endogenous *pfrkip* gene was appended with a V5-epitope tag at the 3’ end using CRISPR-Cas9 (Fig. 2). A single plasmid system was created with Cas9 expression cassette along with a guide DNA sequence specific to the 3’ end of *pfrkip* gene (Fig. 2a). The cloning of guide was verified by DNA sequencing (not shown). Clonal parasites containing V5 epitope tag were verified through a diagnostic PCR (Fig. 2b). Primer pair F4/R3 that amplifies the entire gene sequence resulted in a specific amplicon of desired size of 1046 bp only in the clonal V5-tagged parasites (Fig. 2b). Another primer pair, F5/R2 selected to amplify specifically in the clonal parasites resulted in an amplicon of 660 bp only in the clonal parasites and not in the WT parasite (Fig. 2b). Contamination of WT parasite in the clonal parasites was ruled out using the primer pair F3/R2. The primer pair F3/R2 resulted in an amplicon of desired size (525 bp) only in the WT parasites (Fig. 2b). To further confirm the desired gene editing, the entire *pfrkip* gene was amplified with F4/R2 and sequenced using internal primers. The presence of V5 epitope tag separated by a small linker sequence and in frame with *pfrkip* gene was confirmed by the chromatogram (Fig. 2c). These results confirm successful tagging of endogenous *pfrkip* gene with a V5 epitope tag at the 3’ end. One of the clones of *Pf:rkipv5*, clone C4 was used to test the expression of chimeric PfRKIP-V5 protein. Lysates prepared from the schizont stage of WT and *Pf*:*rkipv5c4* were separated on SDS-PAGE followed by Western blotting using anti-V5 antibodies. A specific band corresponding to the size of chimeric PfRKIP-V5 was detected specifically in the *Pf*:*rkipv5c4* parasites (Fig. 2d). Two parallel blots were probed with anti-PfRKIP and anti-β-actin antibodies to confirm the presence of PfRKIP and similar loading in the WT and *Pf*:*rkipv5c4* parasites (Fig. 2d). Taken together, these results confirm the expression of PfRKIP in the asexual blood stages of the parasite that peaks during the late schizogony.

**Figure 1.**
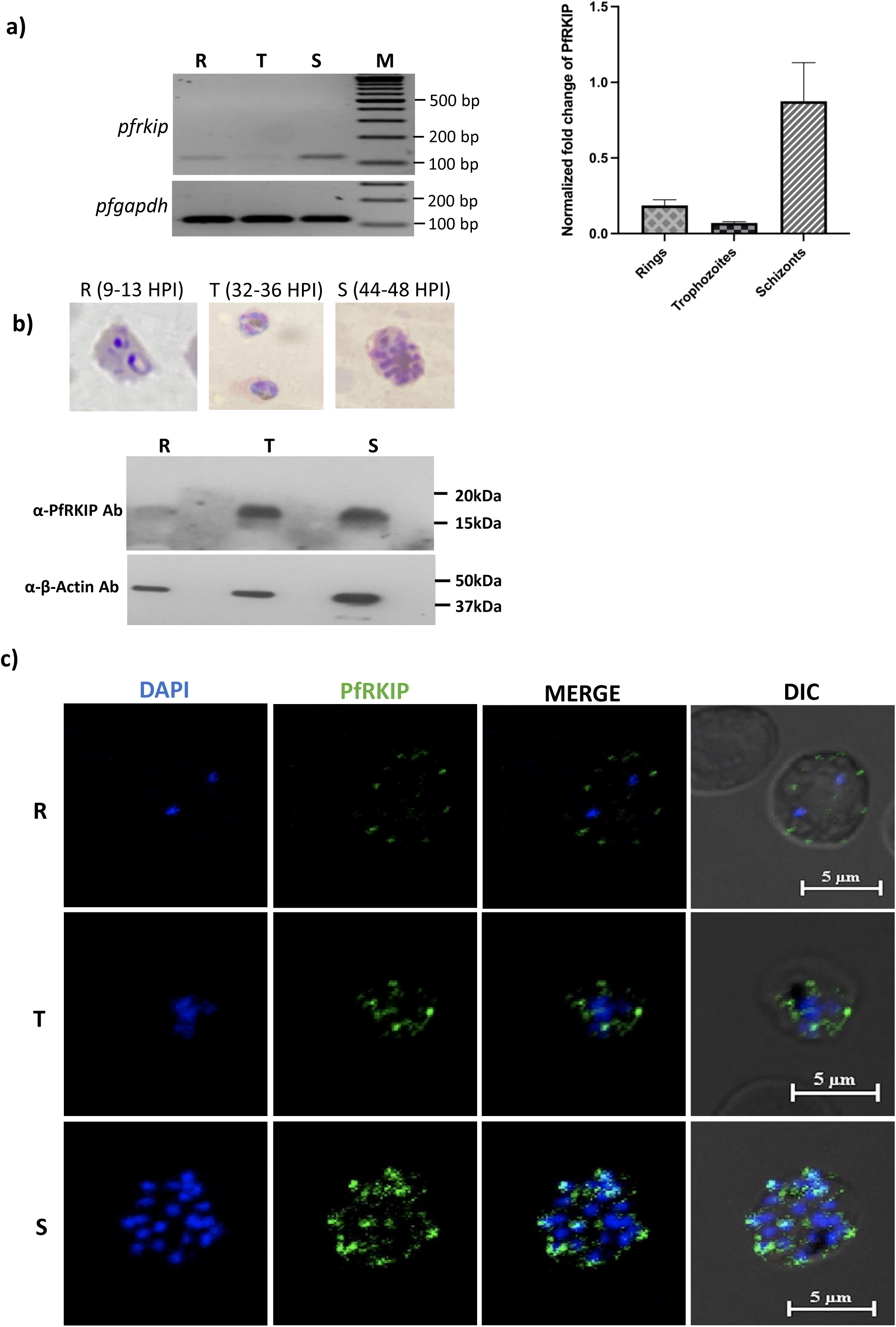
PfRKIP is expressed throughout the asexual blood stages of *P*. *falciparum*. **a)** A semiquantitative RT-PCR was carried out with cDNA prepared from highly synchronized ring (R), trophozoite (T) and schizont (S) stage parasite and separated on agarose gel. An amplicon of 120 bp specific for *pfrkip* is detected. *pfgapdh* was used as a loading control. M-GeneRuler DNA Ladder Mix. The expression of *pfrkip* in each stage of the parasite was normalized with *pfgapdh* expression for the same stage and plotted as a histogram from two independent biological experiments. Normalized fold change of *pfrkip* on the Y-axis is plotted against ring, trophozoite and schizont on the X-axis. Error bars represent standard error of mean. **b)** PfRKIP protein is expressed in all the asexual stages of the parasite. Parasite lysates prepared from R, T and S were separated on SDS-PAGE and probed with anti-PfRKIP antisera through a Western blot. A parallel blot was probed with anti-beta actin antibodies (α-β-actin Ab) as a loading control. A representative blot from two independently performed experiments is shown. **c)**. Immunofluorescence assay shows expression of PfRKIP in asexual blood stages. PfRKIP was detected in R, T and S stage of *P*. *falciparum* by IFA using anti-PfRKIP antibodies and Alexa Fluor 488-conjugated goat anti-rat IgG (green). Nuclear DNA was counterstained with 4′,6-diamidino-2-phenylindole (DAPI, blue). DIC-Differential Interference Contrast. White scale bar is 5 μm.

**Figure 2.**
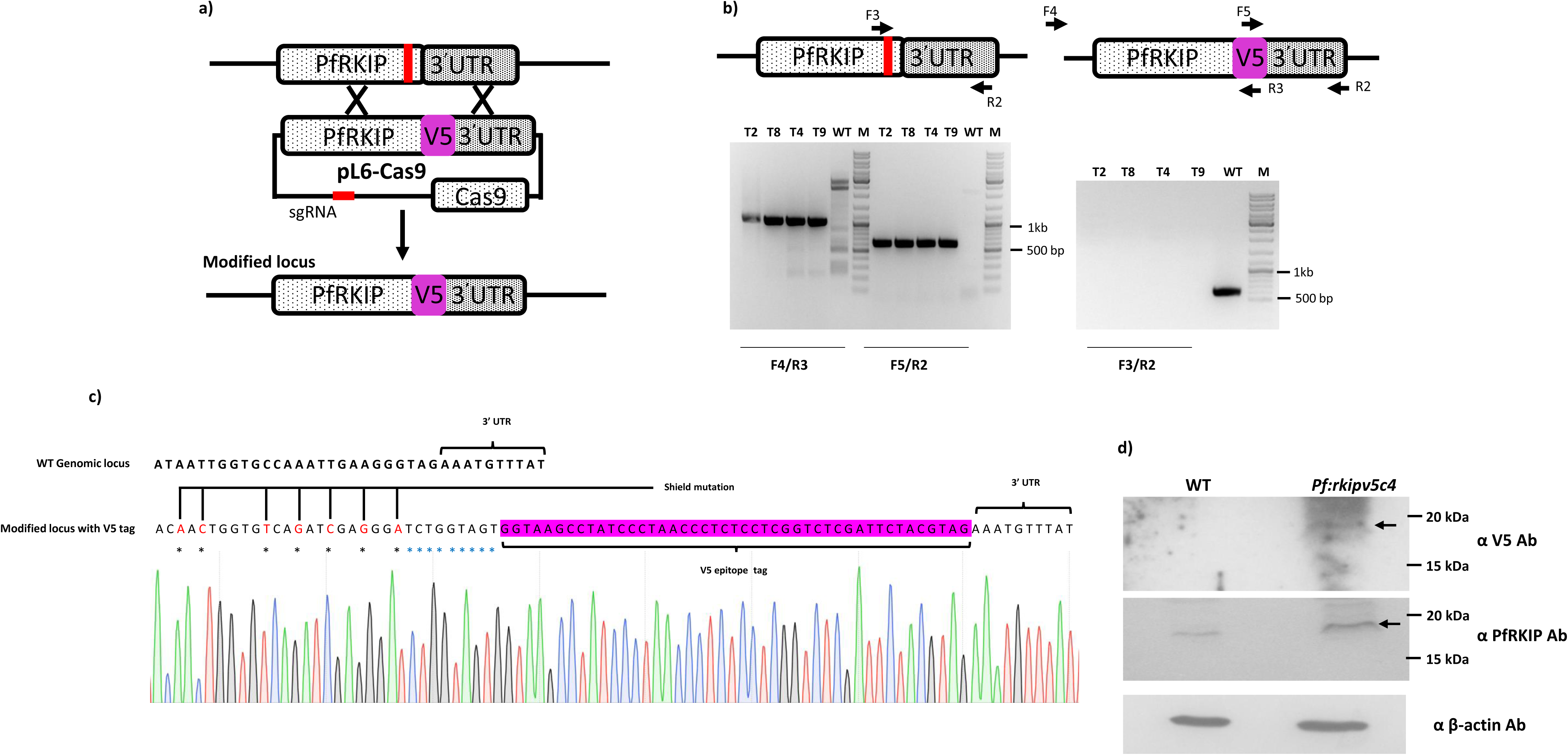
Epitope tagging of endogenous *pfrkip* through CRISPR-Cas9. **a)** Schematic representation of the strategy used for tagging the endogenous *pfrkip* gene at the 3’ end with V5 epitope tag. The region of *pfrkip* targeted for double strand break by Cas9 endonuclease is represented by a red solid line. A sgRNA containing the guide DNA sequence corresponding to the targeted *pfrkip* gene sequence was cloned in pL6-Cas9 plasmid. The modified *pfrkip* locus is represented after the desired gene editing. **b)** Diagnostic PCR for verification of site-specific integration of V5 epitope tag. Positions of the oligonucleotides used for diagnostic PCR are shown in the schematic architecture of *pfrkip*. Two primer pairs, F4/R3 and F5/R2 selected for specific amplification in V5 tagged transgenic parasites, result in amplicons of desired sizes, 1046 bp and 660 bp, respectively in the clonal parasites (T2, T4, T8, T9). Primer pair F3/R2, specific for WT sequence, result in an amplicon of 525 bp only in the WT parasites while no amplification was observed in the V5 tagged clonal parasites. **c)** Verification of endogenous *pfrkip* locus for V5 epitope tagging through DNA sequencing. The entire gene sequence from the modified transgenic parasites was PCR amplified using F4/R2 primer pair and sequenced through Sanger DNA sequencing using internal primers. Chromatogram of a selected region depicting V5 epitope tag (pink) and linker sequence (blue asterisk) are represented. Shield mutations incorporated to avoid repeated DNA cutting by Cas9 are shown as black asterisk. **d)** V5 epitope tagged PfRKIP is detected in the schizonts of *Pf:rkipv5*. Expression of V5 tagged PfRKIP was detected in the schizonts of *Pf:rkipv5* clone 4 using anti-V5 antibody. PfRKIPV5 of desired size (~ 23 kDa) was observed specifically in the C4 clone (black arrow) whereas no band at the corresponding position was detected in the WT parasite. A parallel blot was probed with anti-PfRKIP and anti-Beta-Actin Antibody (α β-actin Ab). The band corresponding to PfRKIP-V5 in the V5 tagged C4 clone is slightly higher compared to that in the WT parasite. A representative blot from two independently performed experiments is shown.

We also tested the localization of PfRKIP in different blood stages of the parasite through immunofluorescence assay (IFA). RBCs infected with ring, trophozoite and schizont stage of the parasite were smeared on a glass slide and processed for IFA. PfRKIP was detected in ring, trophozoite and schizont stage of the WT parasite (Fig. 1c). PfRKIP was localized as discrete dots within the parasite in ring and trophozoite (Fig. 1c). In a mature schizont, PfRKIP was seen as distinct foci with some level of spilling around the foci (Fig. 1c). In highly mature segmented schizont stage, the spreading around the foci was reduced and discrete puncta associated with the developing merozoites were observed (Fig. 1c and Fig. 8). We also attempted to localize PfRKIP in the *Pf*:*rkipv5* parasites using anti-V5 antibodies, however due to non-specific staining in the WT parasites with anti-V5 antibodies no conclusive result could be drawn (result not shown). Nevertheless, staining of *Pf*:*rkipv5c4* schizonts with anti-PfRKIP antibodies result in a similar staining pattern as in the WT parasites (Fig. 8b). Furthermore, we also tested the localization of PfRKIP in mature gametocytes (stage IV and V) through IFA. In stage IV and V gametocytes, PfRKIP showed peripheral staining pattern suggesting preferential localization in the parasite membrane or parasitophorous vacuole membrane that could perhaps be due to its interaction with the phospholipids. Additionally, in male stage V gametocyte PfRKIP was also found localized within the cytoplasm (Fig. S2, top panel). In female stage V gametocyte, we noticed higher intensity with anti-PfRKIP Ab compared to that in male stage V gametocyte suggesting that PfRKIP expression may be more in female gametocytes versus male (Fig. S2). Interestingly, in female stage V gametocyte, PfRKIP was also found associated with the RBC membrane that was not evident in stage IV and male stage V gametocytes (Fig. S2). Taken together, these results show that PfRKIP is present in all the asexual blood stages and the punctate staining pattern observed in segmented schizonts suggest that PfRKIP may be involved in processes required for egress and/or invasion of RBCs. In addition, PfRKIP is also present in mature gametocytes where it may have a role during sexual development of the parasite.

### PfRKIP show pH dependent distinct interaction profile with lipids

PEBP domain containing proteins are known to interact with lipids including phosphatidyl ethanolamine [16–18]. PfRKIP contains a putative PEBP domain with well conserved signature motif (DPDXP) unique to the PEBP family [16](Fig. 3a). Apart from two regions within PEBP domain with high sequence conservation (76-80 and 124-130), the PfRKIP protein sequence shows very less overall conservation compared to its human counterpart, HsRKIP (Fig. 3a). The N- and C-terminus of the proteins are highly variable in primary amino acid sequence (Fig. 3a) and have been shown in other PEBP domain containing proteins to determine the crucial structural features required for recognition of cognate ligands [16, 19, 20]. We were interested in testing the interaction of PfRKIP with different physiologically relevant lipids and compare the lipid binding profile with HsRKIP. To this end, we expressed the complete CDS of both PfRKIP and HsRKIP in *E*. *coli* and purified the two proteins to homogeneity using metal affinity chromatography (Fig. 3b). The full length PfRKIP and HsRKIP migrated at the expected molecular weight of ~ 23 kDa on SDS-PAGE (Fig. 3bi and 3bii, respectively). The identity of the two recombinant proteins was verified through Western blot using anti-HIS tag antibodies since both the proteins were expressed with a C-terminal 8X HIS-tag. Bands with the desired molecular weights (~ 23 kDa), corresponding to PfRKIP and HsRKIP on the SDS-PAGE were detected in the Western blot with anti-HIS antibodies (Fig. 3b). The identity of both the proteins was also verified by mass spectrometry (data not shown).

**Figure 3.**
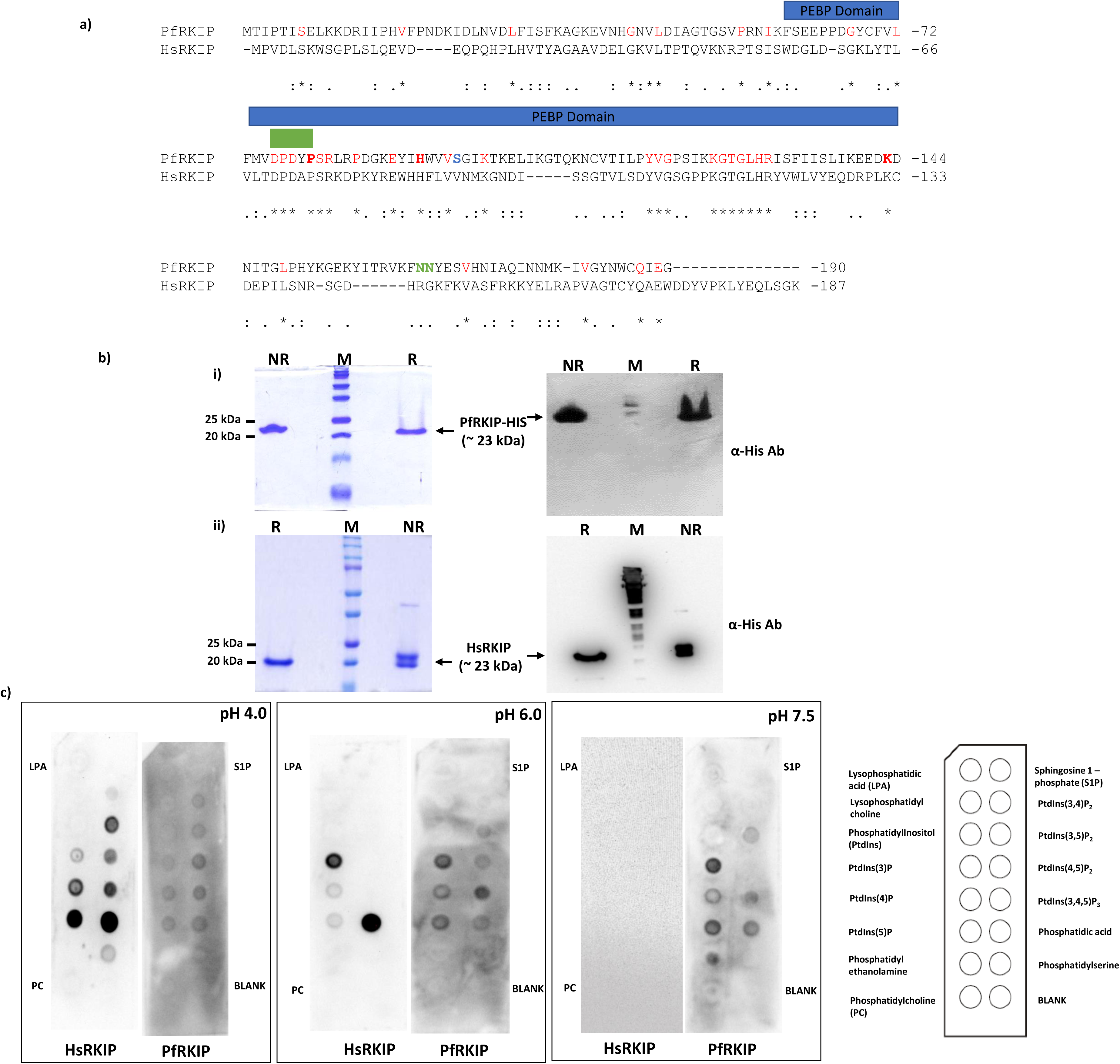
Recombinant PfRKIP shows pH dependent, distinct lipid interaction profile. **a)** Structure based sequence alignment of PfRKIP and *Homo sapiens* RKIP (HsRKIP) [65]. PfRKIP contains a putative phosphatidylethanolamine binding protein (PEBP) domain (blue box) that consist of a region with signature motif (DPDXP) (green box). Amino acid residues highlighted in bold red, blue and green were selected for further biochemical studies. Red: conserved amino acids; Blue: phosphorylation site under *in vitro* conditions; Green: part of the left-handed alpha helix (α5) that is absent in HsRKIP. **b)** Recombinant PfRKIP and HsRKIP proteins were purified to homogeneity. PfRKIP and HsRKIP were expressed with a C-terminal 8X histidine tag and purified to homogeneity with Ni-NTA sepharose beads using metal affinity chromatography. The recombinant proteins were separated on 12 % SDS-PAGE under non-reducing and reducing conditions. Parallel SDS-PAGE gels were used for Western blot and probed with anti-HIS antibody (α-His Ab). Bands corresponding to recombinant PfRKIP and HsRKIP on the SDS-PAGE were detected with α-His Ab. M: pre-stained molecular weight standard. **c)** Recombinant PfRKIP shows pH dependent distinct interaction profiles with lipids. A commercially available membrane containing 15 immobilized lipids (shown as inset on the right) was incubated with recombinant PfRKIP and HsRKIP at three different pH conditions (pH: 4.0, 6.0 and 7.5). At different pH conditions, PfRKIP and HsRKIP show distinct interaction profiles with the immobilized lipids.

The recombinant PfRKIP and HsRKIP were used to test their interaction with different lipids through a far-Western approach. A commercially available hydrophobic membrane strip containing 15 different immobilised lipids was incubated with 20 μg/mL of recombinant PfRKIP or HsRKIP. The bound recombinant proteins were detected with anti-HIS antibodies. Interaction of recombinant proteins with the lipids was tested at three different pH conditions: pH 4.0, 6.0 and 7.5. Interaction of PfRKIP with monophosphorylated phosphatidylinositols: phosphatidylinositol-3-phosphate (PtdIns(3)P), phosphatidylinositol-4-phosphate (PtdIns(4)P), and phosphatidylinositol-5-phosphate (PtdIns(5)P) increased from pH 4.0 to pH 7.5 (Fig. 3c). At pH 4.0 and pH 6.0, PfRKIP was found to interact with all the three phosphatidylinositol bisphosphates: PtdIns(3,4)P_2_, PtdIns(3,5)P_2_, and PtdIns(4,5)P_2_ (Fig. 3c). However, at pH 7.5, PfRKIP showed interaction with only PtdIns(3,5)P_2_ form of phosphatidylinositol bisphosphates (Fig. 3c). Interaction of PfRKIP with PtdIns(3,4,5)P_3_, the trisphosphorylated form of phosphatidylinositol, and phosphatidic acid was observed under all the pH conditions tested (Fig. 3c). Interestingly, interaction of PfRKIP with phosphatidylethanolamine was evident only at the physiological pH (Fig. 3c). No interaction of PfRKIP was observed with phosphatidylinositol (PtdIns) at any pH condition suggesting that PfRKIP binds to PtdIns only when it is phosphorylated at specific sites (Fig. 3c). In addition, PfRKIP did not show any perceptible interaction with sphingosine1-phosphate, lysophosphatidic acid, lysophosphatidyl choline, phosphatidylserine and phosphatidylcholine at any pH condition (Fig. 3c). Importantly, HsRKIP showed distinct interaction profile with lipids compared to PfRKIP. A strong interaction was observed between HsRKIP and phosphatidic acid at pH 4.0 and pH 6.0 compared to PfRKIP. At pH 4.0, the interaction of HsRKIP with phosphorylated phosphatidylinositols was more compared to PfRKIP at the same pH. Overall the interaction of HsRKIP with lipids decreased with increase in pH. Surprisingly, at pH 7.5, no interaction was observed between HsRKIP and any of the lipids under the same conditions as PfRKIP suggesting that perhaps HsRKIP requires higher concentration of lipids or specific cellular environment for recognition of specific lipid molecules. Taken together, our results show that PfRKIP exhibit differential interaction profiles at different pH conditions that are distinct from its human counterpart. It may be possible that specific modifications of lipid molecules by cellular machinery under various environmental cues may play an important role in determining the activation of specific signaling cascades through PfRKIP [21, 22].

In order to gain more insight into the specific recognition mechanism between PfRKIP/HsRKIP and lipids at different pH conditions, we employed molecular docking approaches using bioinformatics. Crystal structures of PfRKIP (PDB ID: 2R77, pH 5.5), and HsRKIP (PDB ID:1BD9, pH 6.5) were used to dock 6 different polar groups associated with phospholipids. In addition, we also modelled PfRKIP using PvRKIP (PDB ID: 2GZQ, pH 7.5) as a template to compare interaction with lipids groups at pH 7.5. The 6 polar groups utilized for our analysis are Phosphoryl Ethanolamine (PubChem CID - 129735059), Phosphotidylinositol 3,4,5-trisphosphate (PubChem CID - 53477782), Phosphatidic acid (PubChem CID - 446066), and Phosphate (PubChem CID - 1061); phosphatidylinositol 3-phosphate and phosphatidylinositol 5-phosphate were drawn using MarvinSketch tool (see the materials and methods; [23]. The binding energy of phosphatidylinositol 3-phosphate and phosphatidylinositol 5-phosphate with PfRKIP was low at pH 7.5 (−8.75 kJ/mol, −8.86 kJ/mol) compared to PfRKIP at pH 5.5 (−8.63 kJ/mol, −8.49 kJ/mol) and HsRKIP at pH 6.5 (−5.04 kJ/mol, −6.21 kJ/mol (Table 1). However, the binding energy of phosphatidylinositol 3,4,5-trisphosphate with PfRKIP was low at pH 5.5 (−10.70 kJ/mol) compared to pH 7.5 (−8.05 kJ/mol) and HsRKIP at pH 6.5 (−6.79 kJ/mol) (Table 1). These results are in agreement with the lipid binding assays performed with the recombinant PfRKIP at different pH conditions (Fig. 3c). Furthermore, in concurrence with the *in vitro* lipid binding assays, HsRKIP at pH 6.5 showed lowest binding energy (−5.68 kJ/mol) with phosphatidic acid compared to that with PfRKIP at pH 5.5 (−3.49 kJ/mol) and pH 7.5 (−3.10 kJ/mol) (Table 1). Like phosphatidylinositol 3,4,5-trisphosphate, phosphorylethanolamine and phosphate show high affinity and low binding energy for PfRKIP at pH 5.5 (−5.75 kJ/mol, −5.12kJ/mol) compared to PfRKIP at pH 7.5 (−4.79 kJ/mol, −4.69 kJ/mol) and HsRKIP at pH 6.5 (−3.20 kJ/mol, −2.54 kJ/mol) (Table 1).

**Table 1.**
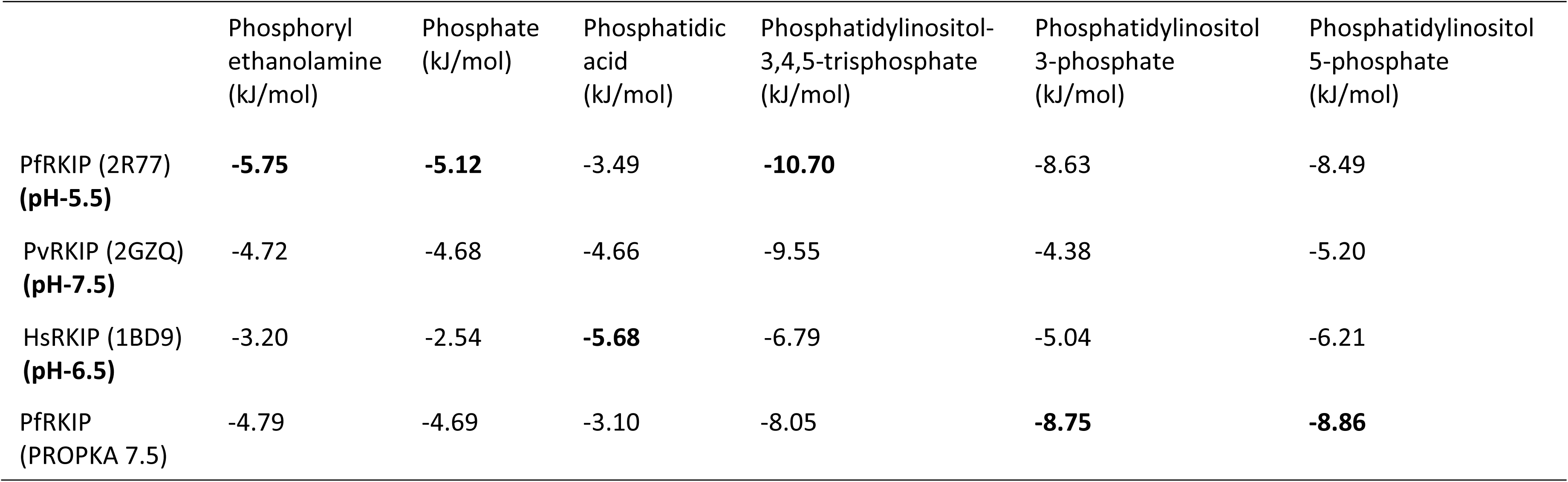
Binding energies (kJ/mol) of various ligands with Proteins.

|  | Phosphoryl<br>ethanolamine<br>(kJ/mol) | Phosphate<br>(kJ/mol) | Phosphatidic<br>acid<br>(kJ/mol) | Phosphatidylinositol-3,4,5-trisphosphate<br>(kJ/mol) | Phosphatidylinositol<br>3-phosphate<br>(kJ/mol) | Phosphatidylinositol<br>5-phosphate<br>(kJ/mol) |
| --- | --- | --- | --- | --- | --- | --- |
| PfRKIP (2R77)<br>**(pH-5.5)** | **-5.75** | **-5.12** | -3.49 | **-10.70** | -8.63 | -8.49 |
| PvRKIP (2GZQ)<br>**(pH-7.5)** | -4.72 | -4.68 | -4.66 | -9.55 | -4.38 | -5.20 |
| HsRKIP (1BD9)<br>**(pH-6.5)** | -3.20 | -2.54 | **-5.68** | -6.79 | -5.04 | -6.21 |
| PfRKIP<br>(PROPKA 7.5) | -4.79 | -4.69 | -3.10 | -8.05 | **-8.75** | **-8.86** |

All the ligands except phosphatidic acid form hydrogen bonding with Lys 123 and Lys 124 of PfRKIP at pH 5.5 and pH 7.5 (Table 2, Fig. 4, Fig. S3a and S3b). While Tyr 120 of HsRKIP interact with ligands and form hydrogen bond except phosphatidyl inositol-3,4,5-triphosphate (Table 2, Fig. 4, Fig. S3c). RKIP of *P*. *vivax* (PvRKIP) another human malaria parasite and a close homolog of *P*. *falciparum*, show involvement of Arg 83 in formation of hydrogen bonds with all the six ligands (Table 2). This suggests that the amino acid residues: Lys 123 and Lys 124 in PfRKIP, Arg 83 in PvRKIP, and Tyr 120 in HsRKIP may be crucial in maintaining the interaction of RKIP with lipids.

**Figure 4.**
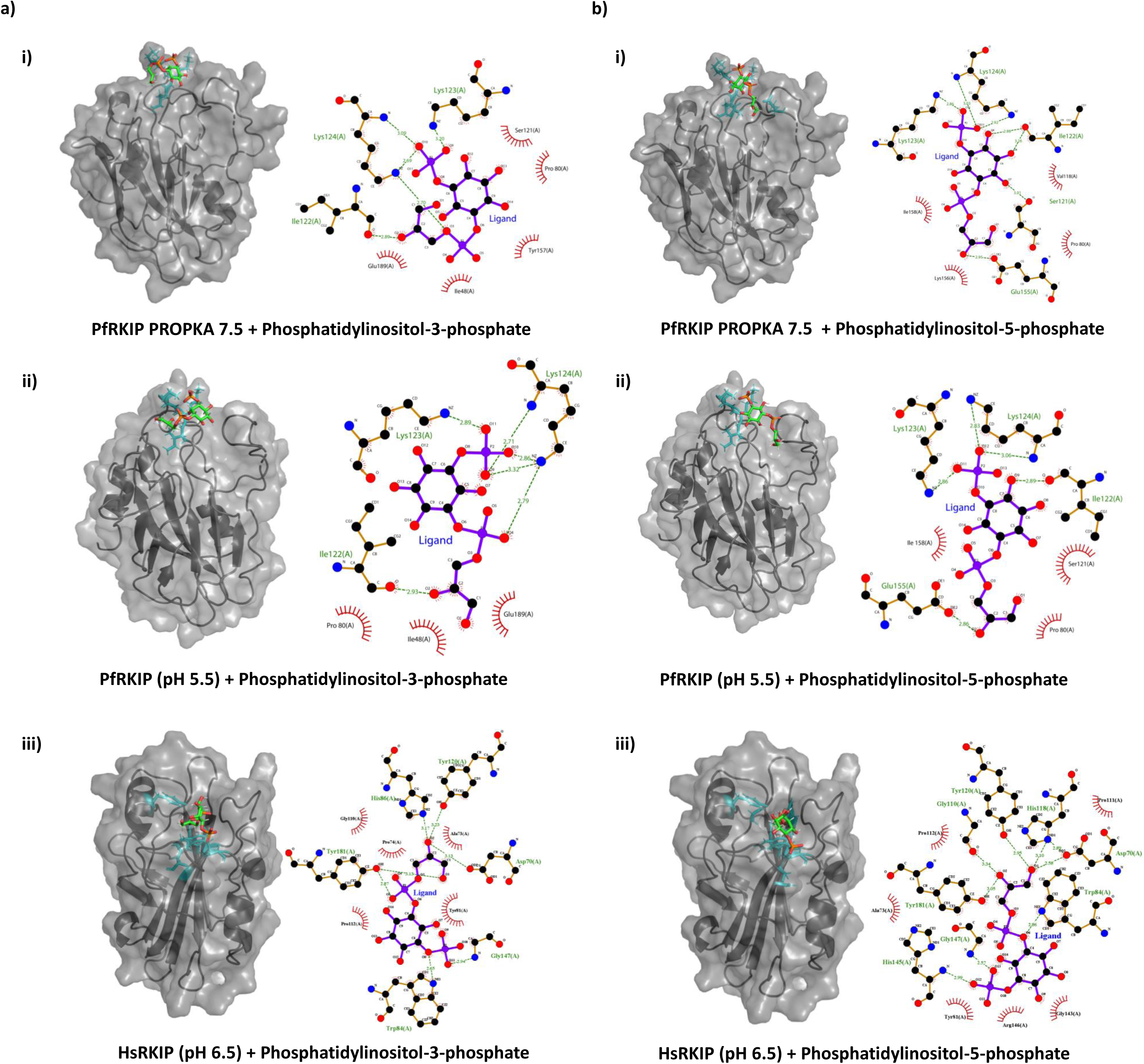
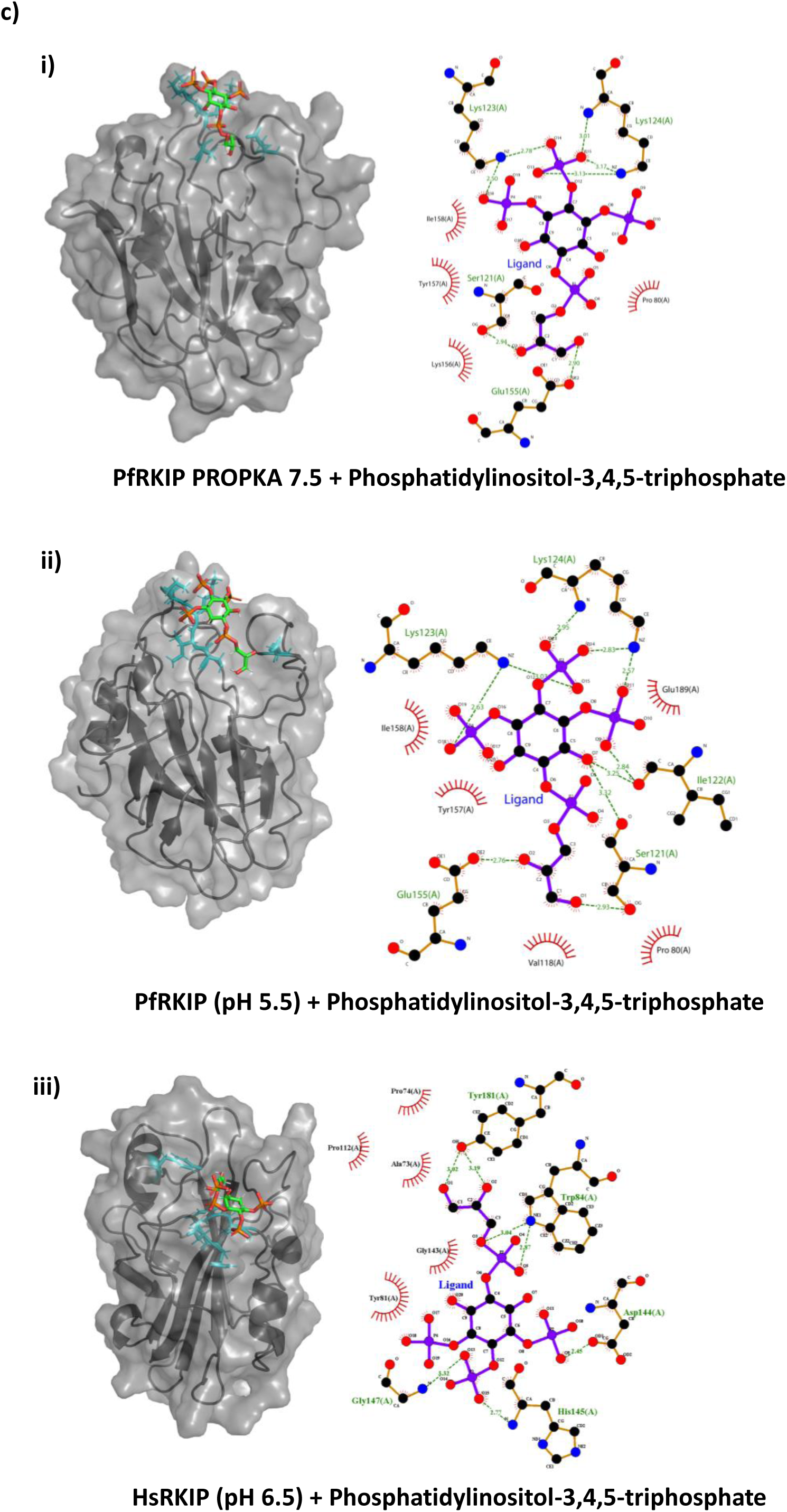
Interaction of phosphatidylinositols with PfRKIP and HsRKIP. Monophosphorylated phosphatidylinositols: phosphatidylinositol-3-phosphate **(a)**, phosphatidylinositol-5-phosphate **(b)** and triphosphorylated phosphatidylinositol: phosphatidylinositol-3,4,5-trisphosphate **(c)** were docked with **i)** PfRKIP (PDB ID: 2R77) modelled at pH 7.5 (PfRKIP PROPKA 7.5), **ii)** PfRKIP at pH 5.5 (PDB ID: 2R77) and **iii)** *Homo sapiens* RKIP (HsRKIP) at pH 6.5 (PDB ID: 1BD9). The 3D structures of corresponding proteins docked with ligands (orange and green sticks) and LigPlot are shown. The interacting residues in the 3D model are represented by cyan color.

**Table 2.**
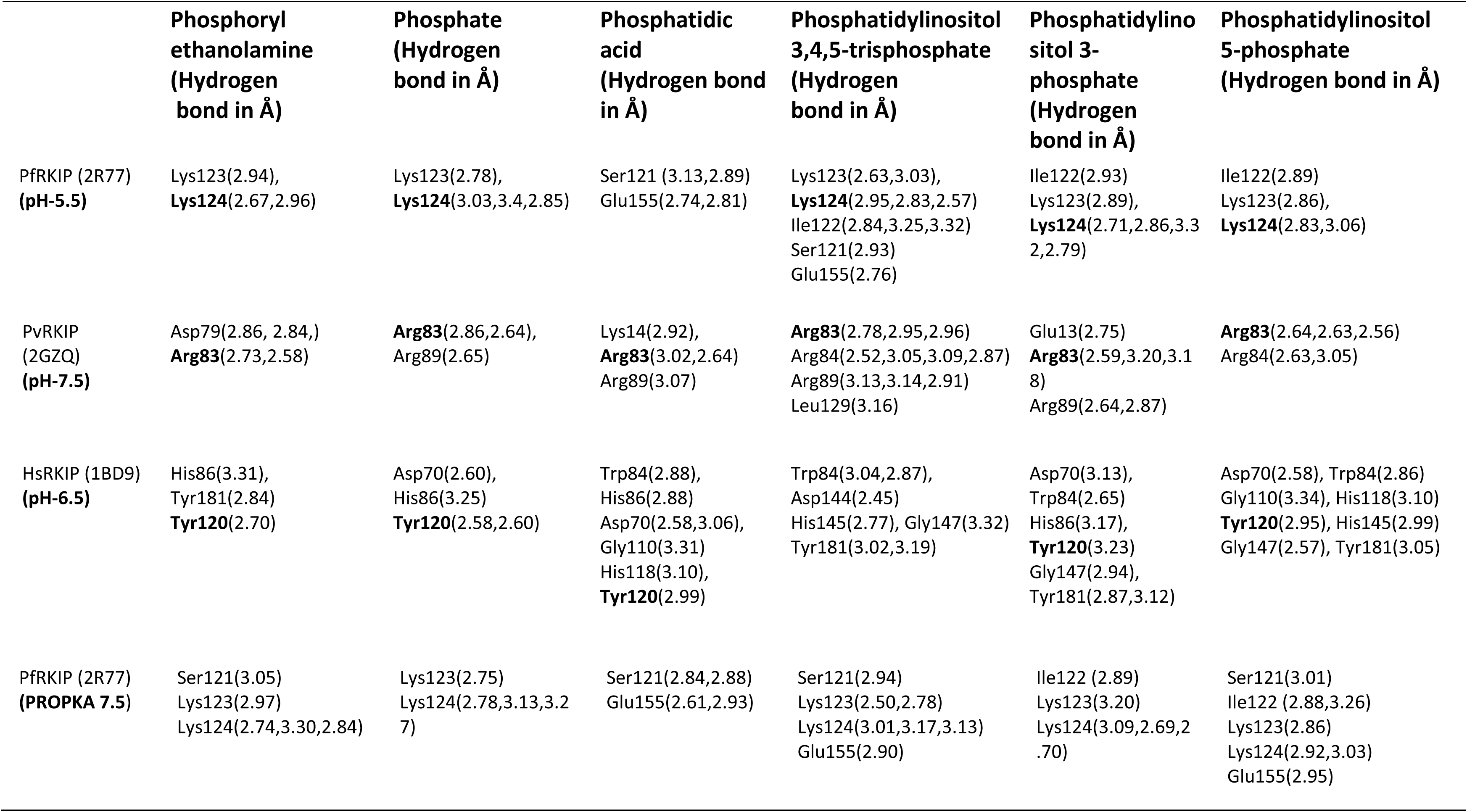
Hydrogen-bond interactions between amino acids of each protein with the corresponding docked ligand.

|  | Phosphoryl<br>ethanolamine<br>(Hydrogen<br>bond in Å) | Phosphate<br>(Hydrogen<br>bond in Å) | Phosphatidic<br>acid<br>(Hydrogen bond<br>in Å) | Phosphatidylinositol<br>3,4,5-trisphosphate<br>(Hydrogen<br>bond in Å) | Phosphatidylinositol 3-phosphate<br>(Hydrogen<br>bond in Å) | Phosphatidylinositol<br>5-phosphate<br>(Hydrogen bond in Å) |
| --- | --- | --- | --- | --- | --- | --- |
| PfRKIP (2R77)<br>(pH-5.5) | Lys123(2.94),<br>**Lys124**(2.67,2.96) | Lys123(2.78),<br>**Lys124**(3.03,3.4,2.85) | Ser121 (3.13,2.89)<br>Glu155(2.74,2.81) | Lys123(2.63,3.03),<br>**Lys124**(2.95,2.83,2.57)<br>Ile122(2.84,3.25,3.32)<br>Ser121(2.93)<br>Glu155(2.76) | Ile122(2.93)<br>Lys123(2.89),<br>**Lys124**(2.71,2.86,3.3<br>2,2.79) | Ile122(2.89)<br>Lys123(2.86),<br>**Lys124**(2.83,3.06) |
| PvRKIP<br>(2GZQ)<br>(pH-7.5) | Asp79(2.86, 2.84,)<br>**Arg83**(2.73,2.58) | **Arg83**(2.86,2.64),<br>Arg89(2.65) | Lys14(2.92),<br>**Arg83**(3.02,2.64)<br>Arg89(3.07) | **Arg83**(2.78,2.95,2.96)<br>Arg84(2.52,3.05,3.09,2.87)<br>Arg89(3.13,3.14,2.91)<br>Leu129(3.16) | Glu13(2.75)<br>**Arg83**(2.59,3.20,3.1<br>8)<br>Arg89(2.64,2.87) | **Arg83**(2.64,2.63,2.56)<br>Arg84(2.63,3.05) |
| HsRKIP (1BD9)<br>(pH-6.5) | His86(3.31),<br>Tyr181(2.84)<br>**Tyr120**(2.70) | Asp70(2.60),<br>His86(3.25)<br>**Tyr120**(2.58,2.60) | Trp84(2.88),<br>His86(2.88)<br>Asp70(2.58,3.06),<br>Gly110(3.31)<br>His118(3.10),<br>**Tyr120**(2.99) | Trp84(3.04,2.87),<br>Asp144(2.45)<br>His145(2.77), Gly147(3.32)<br>Tyr181(3.02,3.19) | Asp70(3.13),<br>Trp84(2.65)<br>His86(3.17),<br>**Tyr120**(3.23)<br>Gly147(2.94),<br>Tyr181(2.87,3.12) | Asp70(2.58), Trp84(2.86)<br>Gly110(3.34), His118(3.10)<br>**Tyr120**(2.95), His145(2.99)<br>Gly147(2.57), Tyr181(3.05) |
| PfRKIP (2R77)<br>(PROPKA 7.5) | Ser121(3.05)<br>Lys123(2.97)<br>Lys124(2.74,3.30,2.84) | Lys123(2.75)<br>Lys124(2.78,3.13,3.2<br>7) | Ser121(2.84,2.88)<br>Glu155(2.61,2.93) | Ser121(2.94)<br>Lys123(2.50,2.78)<br>Lys124(3.01,3.17,3.13)<br>Glu155(2.90) | Ile122 (2.89)<br>Lys123(3.20)<br>Lys124(3.09,2.69,2<br>.70) | Ser121(3.01)<br>Ile122 (2.88,3.26)<br>Lys123(2.86)<br>Lys124(2.92,3.03)<br>Glu155(2.95) |

### Conserved residues in PEBP domain of PfRKIP are important for its interaction with lipids

Like other PEBP domain containing proteins, PfRKIP contain motifs that are important for interaction with lipids and other partner proteins [16, 24](Fig. 3a). In order to understand the interaction of PfRKIP with its ligands we selected few residues in PfRKIP for mutagenesis study that are either conserved or reported in the literature. We selected Pro 80 which is part of a highly conserved Asp-Pro-Asp-x-**Pro** motif along with another conserved residue His 92 in the PEBP domain (Fig. 3a). Histidine residue at corresponding position in HsRKIP is important for interaction with other proteins [25]. In *P*. *vivax*, RKIP contains a rare left-handed α-helix, α5 that is present near the ligand binding site and is postulated to have a role during recognition of phospholipids [24]. A similar left-handed α-helix is also present at the corresponding position in PfRKIP that we selected for disruption to study its role in lipid binding. Additionally, we selected Lys 143 for mutagenesis studies as the lysine residue at corresponding position in HsRKIP plays an important role in ligand binding [26]. Under *in vitro* conditions, PfRKIP is phosphorylated by PfCDPK1 at Ser 96 [27]. However, how Ser 96 phosphorylation by PfCDPK1 affects the activity of PfRKIP or its interaction with different ligands is not known. Therefore, in order to understand the significance of Ser 96 phosphorylation on the functional aspect of PfRKIP we selected this residue along with others for the mutagenesis study. The selected residues in PfRKIP were mutated as follows: P80L, H92A, S96A, K143A and Δα5. All the mutants of PfRKIP were cloned and verified for the presence of the desired mutation by DNA sequencing (data not shown). The proteins were expressed and purified to homogeneity using metal affinity chromatography (Fig. 5a). All the mutant PfRKIP recombinant proteins conformed to the desired size (~ 23 kDa) and migrated similarly as the WT PfRKIP on SDS-PAGE (Fig. 5a). All the mutant proteins were detected by anti-PfRKIP specific antibodies (Fig. 5a) confirming the identity of the proteins. For thermodynamic characterization of the interaction between the lipid and the recombinant proteins, we employed isothermal calorimetry (ITC). The binding thermogram of WT PfRKIP and the mutant proteins were biphasic in nature except HsRKIP which has monophasic curve (Fig. 5b). The biphasic curve indicates availability of two types of binding sites on the proteins for the lipid. The association of protein with lipid was found to be exothermically driven with loss of entropy. It was observed that binding of wild type and mutant PfRKIP proteins with lipid is quite similar and in the range of ~10^4^ M^-1^ except PfRKIP^P80L^ (Fig. 5b, Table 5). The P80L mutant was found to interact with least affinity with PtdIns(3)P (k_avg_ = 0.6×10^4^) compared to the WT PfRKIP (k_avg_ = 6.5×10^4^). The binding affinity of HsRKIP with PtdIns(3)P, at near physiological pH, was also low (k_avg_ = 1.4×10^4^) compared to that of WT PfRKIP which is in agreement with the lipid binding experiment performed with the recombinant proteins in far-Western format (Fig. 3c). Other PfRKIP mutants: PfRKIP^H92A^, PfRKIP^S96A^, and PfRKIP^Δα5^ also showed decreased binding affinities with PtdIns(3)P (k_avg_ = 5.3×10^4^, 5.5×10^4^, and 5.2×10^4^) although more than that of the P80L mutant (Fig. 5b). These results suggest that the mutations introduced at the selected positions in PfRKIP are important determinants in defining the interaction of the polar head group of PtdIns(3)P with PfRKIP. The PfRKIP^P80L^ mutant exhibited the most dramatic decrease in the binding interaction suggesting that the conserved P80 position in the ligand binding domain is critical for the interaction of PfRKIP with lipids.

**Figure 5.**
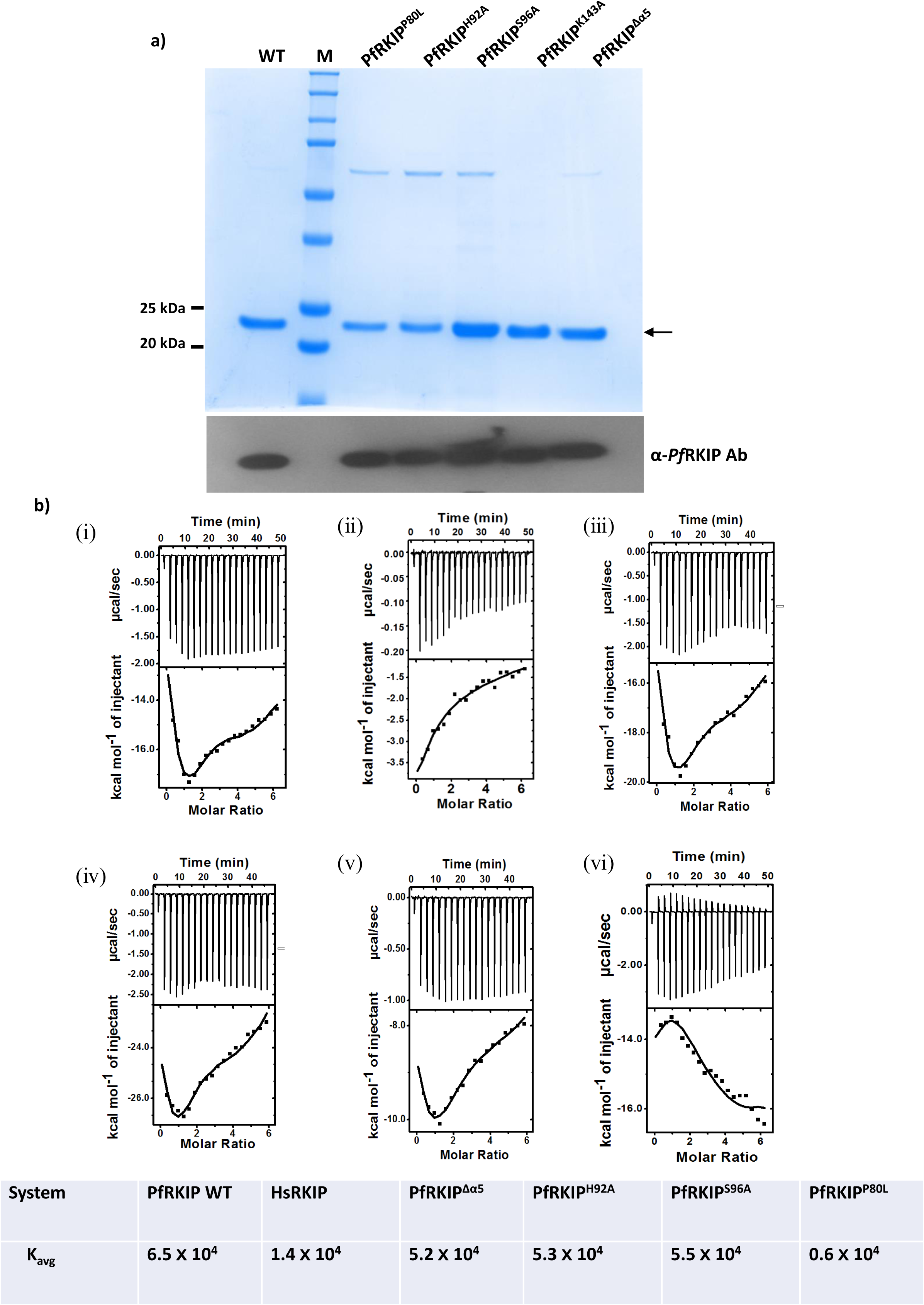
The PEBP domain of PfRKIP contain residues that play important role in lipid recognition. **a)** SDS-PAGE profile of PfRKIP WT and mutant recombinant proteins. Amino acid residues of PfRKIP highlighted in Figure 2 were mutated (P80L, H92A, S96A, K143A, and Δα5) and expressed as recombinant proteins to identify their importance in interaction with PtdIns(3)P. SDS-PAGE stained with Coomassie Blue R250 shows the purified WT and mutant PfRKIP recombinant proteins (black arrow) under reducing condition. A parallel SDS-PAGE gel was used for Western blotting with anti-PfRKIP antibody. **b)** Mutation of residues in the PEBP domain of PfRKIP show reduced interaction with lipid. Recombinant PfRKIP WT and mutant proteins along with HsRKIP were used to test their interaction with PtdIns(3)P through isothermal calorimetry (ITC). ITC thermogram showing the titration of PtdIns(3)P into **(i)** PfRKIP WT; **(ii)** HsRKIP; **(iii)** PfRKIP^Δα5^ **(iv)** PfRKIP^H92A^; **(v)** PfRKIP^S96A^; and **(vi)** PfRKIP^P80L^. The concentration of all proteins and lipid were 20 µM and 600 µM, respectively. All the experiments were performed in phosphate buffer [1.47 mM KH_2_PO_4_, 4.3 mM Na_2_HPO_4_ and 137 mM NaCl] at pH 7.6. PfRKIP^P80L^ shows the greatest reduction in interaction with PtdIns(3)P compared to the other mutant proteins. The value of average binding constant (K_avg_) is tabulated for all the recombinant proteins used in the experiment.

### Recombinant PfRKIP interacts with PfCDPK1

An earlier study suggested that PfRKIP might act as a substrate for PfCDPK1 in the parasite based on *in vitro* kinase experiments with recombinant proteins [27]. However, molecular determinants for the interaction between the two proteins have remained elusive so far. Therefore, in order to understand the molecular mechanism for the recognition /interaction between PfCDPK1 and PfRKIP, we used the PfRKIP mutants and studied their interaction with PfCDPK1 through ELISA. The recombinant PfCDPK1 was coated on a 96 well plate and allowed to incubate with increasing concentrations of different PfRKIP proteins. The binding of PfRKIP protein with PfCDPK1 was detected through anti-PfRKIP antibody. The binding of recombinant WT PfRKIP with PfCDPK1 increased with increasing concentration of PfRKIP that saturated at around 10 μg/mL (Fig. 6a). The binding of PfRKIP with PfCDPK1 was maximally affected in PfRKIP^K143A^ followed by PfRKIP^H92A^ and PfRKIP^P80L^ compared to the WT PfRKIP. The decrease in binding of PfRKIP mutants: PfRKIP^K143A^, PfRKIP^H92A^, and PfRKIP^P80L^ with recombinant PfCDPK1 suggest that K143, H92 and P80 may be important for the interaction of PfRKIP with PfCDPK1. The decrease in interaction of PfRKIP^K143A^, PfRKIP^H92A^, and PfRKIP^P80L^ with PfCDPK1 was found to be statistically significant (p=0.0128, p=0.0023, p=0.002; n=3; two-tailed paired t-test). The PfRKIP^Δα5^ mutant did not show any significant difference in interaction with PfCDPK1 compared to the WT PfRKIP (Fig. 6a) suggesting that the left-handed α-helix, α5 is not an essential determinant for the interaction of PfRKIP and PfCDPK1. The PfRKIP^S96A^ mutant showed decreased interaction with PfCDPK1 compared to the WT suggesting that the loss of a nucleophilic centre at 96 position of PfRKIP, as in S96A leads to decease in its interaction with PfCDPK1. Taken together, these results show that the conserved residues in the PEBP domain of PfRKIP that participate in formation of a ligand binding site are important in interaction of PfRKIP with PfCDPK1.

**Figure 6.**
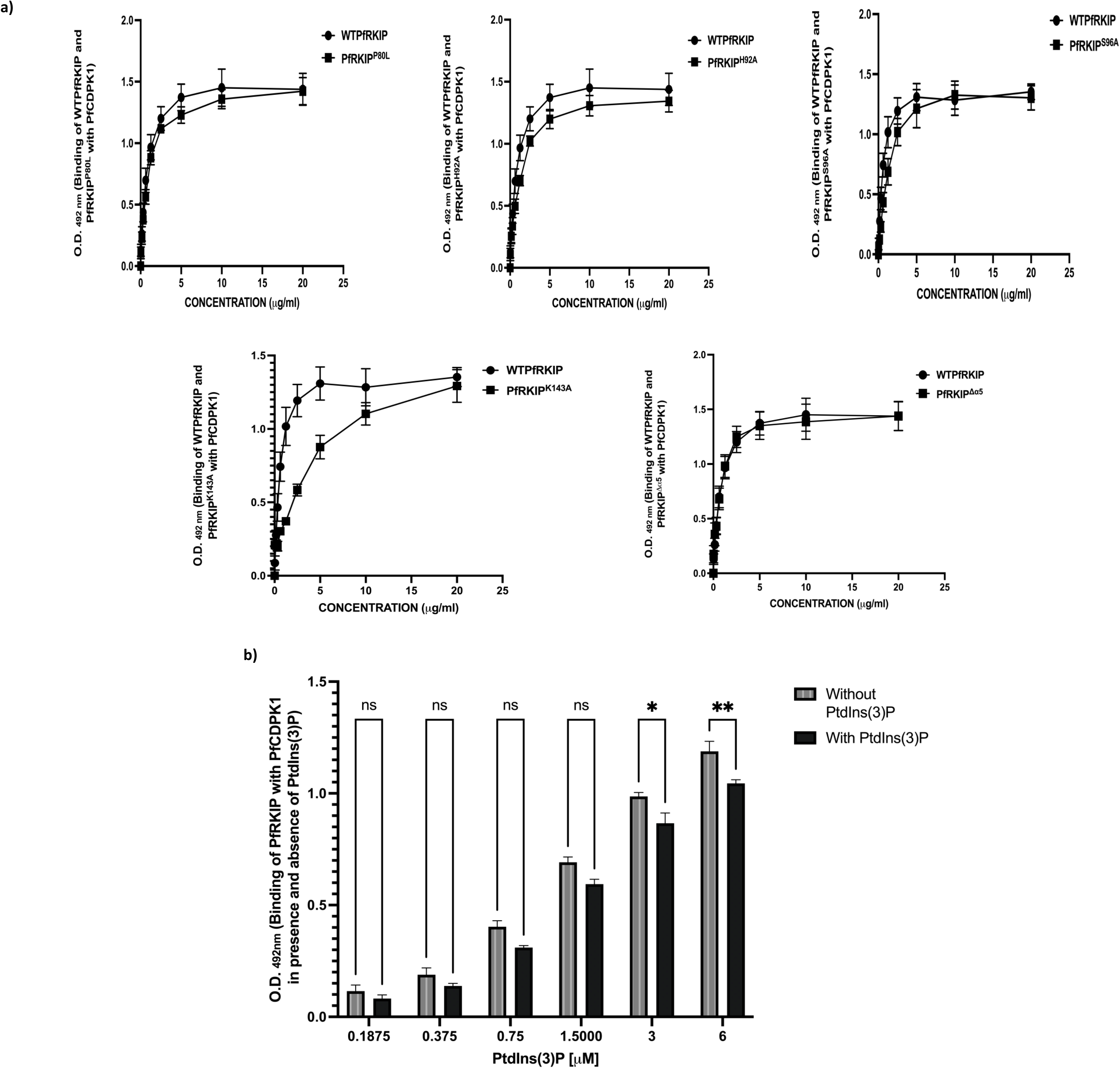
Recombinant PfRKIP interacts with PfCDPK1 and the interaction decreases in presence of a lipid. **a)** Recombinant PfRKIP interacts with recombinant CDPK1. Interaction of recombinant PfRKIP and PfCDPK1 was tested by ELISA. PfCDPK1 protein was coated on the wells of a 96 well plate and incubated with different concentrations of PfRKIP protein. PfRKIP bound with PfCDPK1 was detected through anti-PfRKIP antibody. HRP-conjugated anti-rat antibodies were used to detect the anti-PfRKIP primary antibody followed by addition of a substrate. The plates were read at 492 nm wavelength. PfRKIP^P80L^, PfRKIP^H92A^, PfRKIP^K143A^ and PfRKIP^S96A^ show decrease in interaction with CDPK1 (p=0.002 p=0.0023, p=0.0128, p=0.0101, respectively; paired t-test). Individual graphs depicting O.D. _492 nm_, representing binding of PfRKIP with PfCDPK1 on Y-axis against concentration of PfRKIP in μg/mL on the X-axis were plotted from 3 independent biological experiments performed in duplicate. Error bars represent standard error of mean. **b)** The interaction of PfRKIP with PfCDPK1 decreases in presence of a lipid. PfCDPK1 protein was immobilized on 96 well plate followed by incubation with different concentration of PfRKIP in presence of increasing concentration of PtdIns(3)P. The binding of recombinant PfRKIP with PfCDPK1 was detected using anti-PfRKIP antibodies through ELISA. The binding of PfRKIP with PfCDPK1 was lower in presence of PtdIns(3)P at all the concentrations of PfRKIP and lipid tested compared to without lipid condition. The concentrations of PtdIns(3)P (in μM) on the X-axis are plotted against O.D. _492 nm_, representing binding of PfRKIP with PfCDPK1 in presence and absence of PtdIns(3)P from three independent biological experiments performed in duplicate. The error bars represent standard error of mean from the experiments. The statistical significance was calculated using GraphPad Prism 9 using two-way ANOVA (*p=0.0142 and **p=0.0021, respectively).

### Interaction between PfRKIP and PfCDPK1 decreases in presence of lipids

Our results show that PfRKIP interact with lipids and also with PfCDPK1. We were interested in testing the effect of lipids on the interaction of PfRKIP with PfCDPK1. To this end PfCDPK1 protein was immobilized in 96 well plate followed by incubation with different concentration of PfRKIP in presence of increasing concentration of PtdIns(3)P (Fig. 6b). The binding of recombinant PfRKIP with PfCDPK1 was detected using anti-PfRKIP antibodies in an ELISA based assay as described above. In presence of PtdIns(3)P, the binding of PfRKIP with PfCDPK1 was lower compared to without PtdIns(3)P at all the concentrations of PfRKIP and lipid tested (Fig. 6b). The decrease in binding of PfRKIP with PfCDPK1 in presence of lipid was statistically significant at 3 μM and 6 μM concentration of PtdIns(3)P (Fig. 6b) (p=0.0142 and p=0.0021, respectively; Two-way ANOVA; n=3 performed in duplicate).

### PfRKIP regulates the kinase activity of PfCDPK1

A previous study has shown that PfRKIP is phosphorylated by PfCDPK1 under *in vitro* conditions [27]. PfRKIP was shown to decrease the phosphorylation of an exogenous substrate, casein. Additionally, under *in vitro* conditions, PfRKIP was shown to be phosphorylated at S96 position by PfCDPK1. We were interested in understanding the mechanistic basis of regulation of PfCDPK1 activity by PfRKIP. To this end we tested the effect of PfRKIP on the auto and transphosphorylation potential of PfCDPK1. The autophosphorylation of PfCDPK1 increased significantly in presence of recombinant PfRKIP compared to myeline basic protein (MBP), a known exogenous substrate of PfCDPK1, used in the same molar ratio (Fig. 7a) (103 ± 0.9 % and 34 ± 8.9 %, respectively, *p<0.05, paired t-test). In addition, PfRKIP was also phosphorylated by PfCDPK1 (Fig. 7a). This result suggests that PfRKIP substantially increases the autophosphorylation of PfCDPK1 and itself gets phosphorylated by PfCDPK1. Since S96 was shown earlier through *in vitro* studies to be phosphorylated by PfCDPK1, therefore we tested the effect of S96A substitution on the phosphorylation of PfRKIP. In the same molar ratio, the phosphorylation of mutant PfRKIP^S96A^ was lower compared to the WT PfRKIP (Fig. 7c) suggesting that S96 is indeed phosphorylated under *in vitro* conditions. However, since, PfRKIP^S96A^ was phosphorylated albeit to a lower level compared to the WT PfRKIP therefore, other residue/s of PfRKIP are also phosphorylated by PfCDPK1 in addition to S96 under *in vitro* conditions. In order to understand the mechanism of regulation of PfCDPK1 activity by PfRKIP in presence of a substrate, we set up *in vitro* kinase assays with a combination of PfRKIP and MBP. Interestingly, in the presence of MBP, there was significant increase in the phosphorylation of PfRKIP as compared to without MBP condition (Fig. 7b, lane 2 versus lane 4 and corresponding histogram, *p<0.05, unpaired t-test). Additionally, in presence of PfRKIP, there was an increase in the phosphorylation of MBP that increased with the concentration of PfRKIP (Fig. 7b). However, the increase in MBP phosphorylation in presence of PfRKIP was statistically not significant. Nevertheless, our results show that in presence of MBP, the phosphorylation of PfRKIP by PfCDPK1 is increased that also results in the phosphorylation of MBP.

**Figure 7.**
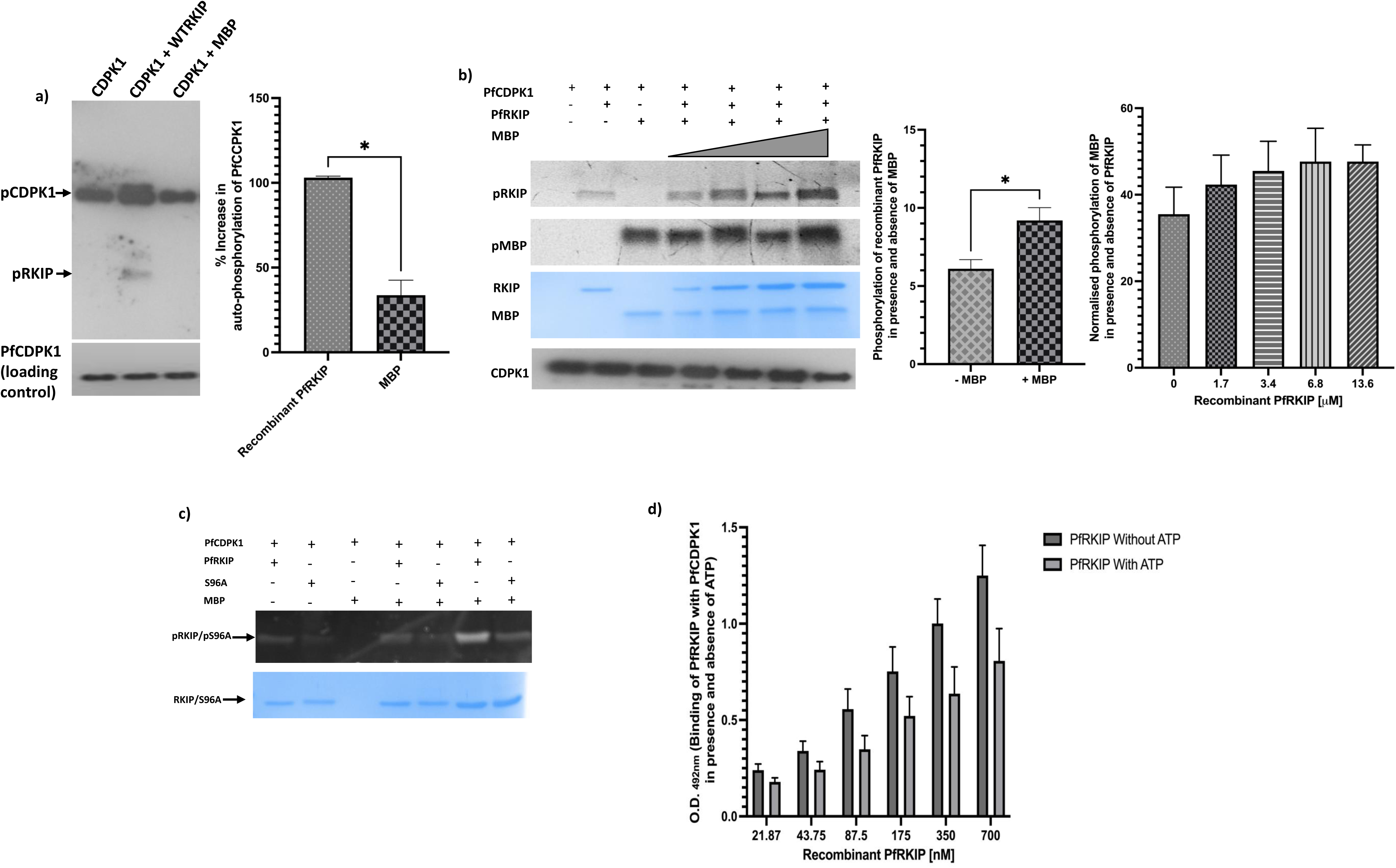
PfRKIP affects the kinase activity of PfCDPK1. **a)** *In vitro* kinase assay with PfCDPK1 was carried out in presence of either recombinant PfRKIP or MBP. Reaction mixtures were separated on SDS-PAGE and processed for Western blotting with anti-thiophosphorylated antibody for the detection of thiophosphorylated PfCDPK1 (pCDPK1). Western blot shows autophosphorylation of PfCDPK1. Presence of PfRKIP leads to significant increase in the autophosphorylation of PfCDPK1. A parallel blot was probed with anti-PfCDPK1 Ab as a loading control. The intensity of autophosphorylated PfCDPK1 was determined using Image J software and histogram was plotted for % increase in autophosphorylation of PfCDPK1 on Y-axis with PfRKIP and MBP on X-axis (n=3, *p<0.05, paired t-test). Error bars represent standard error of mean. **b)** Phosphorylation of PfRKIP is increased in presence of MBP that in turn results in transphosphorylation of MBP. *In vitro* kinase assay was set with different concentrations of PfRKIP in presence of MBP. SDS-PAGE gel stained with ProQ^TM^ diamond show the phosphorylation of PfRKIP (pRKIP) and MBP (pMBP). Phosphorylation of PfRKIP is increased in the presence of MBP (lane 2 versus lane 4, from left to right). A histogram was plotted to represent the phosphorylation of recombinant PfRKIP in presence and absence of MBP (n=3, *p<0.05, unpaired t-test). The phosphorylation of MBP also increases with increasing concentration of PfRKIP in a dose dependent manner. A histogram was plotted with normalized phosphorylation of MBP on the Y-axis against concentration of recombinant PfRKIP (in μM) on the X-axis. The error bars depict standard error of mean. A representative gel image and corresponding histogram from 3 independent biological experiments are shown. **c)** S96A mutation leads to decrease in the phosphorylation of PfRKIP by PfCDPK1. The PfRKIP^S96A^ mutant show less phosphorylation compared to the WT PfRKIP at different concentration used. A representative gel image from two independent biological experiments is shown. **d)** Phosphorylation of PfRKIP decreases its interaction with PfCDPK1. PfCDPK1 was coated on 96 well plate and incubated with increasing concentration of PfRKIP in presence or absence of ATP. The interaction of PfRKIP with PfCDPK1 decreases in the presence of ATP. O.D. _492 nm_, representing binding of PfRKIP with PfCDPK1 in presence and absence of ATP on the Y-axis is plotted against the concentration of PfRKIP on the X-axis. The data is plotted from 3 independent biological experiments performed in duplicate (n=3; p<0.05, paired t-test).

In order to understand the effect of PfRKIP phosphorylation on its interaction with PfCDPK1, we took advantage of an ELISA based assay. PfCDPK1 was coated on a 96 well plate and allowed to incubate with different concentrations of PfRKIP protein either in presence or absence of ATP (100 μM). The bound PfRKIP was detected with anti-PfRKIP Ab. In the presence of ATP, the binding of PfRKIP with PfCDPK1 was lower compared to the absence of ATP at all the concentrations of PfRKIP tested (Fig. 7d). The decrease in the binding of PfRKIP with PfCDPK1 in presence of ATP was statistically significant (p<0.05, n=3, paired t-test). To rule out any adverse effect of immobilization of PfCDPK1 on its kinase activity, we tested the phosphorylation of MBP in the 96 well plate. PfCDPK1 immobilized on the well was indeed catalytically active and was able to phosphorylate MBP in the presence of ATP (Fig. S4). Taken together these results suggest that in presence of PfRKIP the autophosphorylation of PfCDPK1 increases that also results in the phosphorylation of PfRKIP. The phosphorylation of PfRKIP reduces its interaction with PfCDPK1. In presence of MBP, an exogenous substrate, the phosphorylation of PfRKIP by PfCDPK1 is further increased that perhaps is essential for its dissociation from PfCDPK1 and concomitant transphosphorylation of MBP (Fig. 11).

### PfRKIP interacts with PfCDPK1 within the parasites

In order to test whether PfRKIP and PfCDPK1 interact within the parasite, we made use of immunofluorescence microscopy. PfCDPK1 is expressed in the asexual blood stages with maximum expression at the late schizont stage [28–30]. Our IFA results suggest that like PfCDPK1, PfRKIP is present throughout the asexual blood stages with peak expression in the mature schizont stage (Fig. 1). In the late segmented schizont, PfRKIP exhibits punctate staining and is associated with the developing merozoites (Fig. 1c and Fig. 8a). Therefore, we tested the co-localization of PfRKIP and PfCDPK1 in the very late segmented schizonts using anti-PfCDPK1 and anti-PfRKIP specific antibodies. In the segmented schizonts and free merozoites, we obtained similar pattern of staining for PfCDPK1 as reported in previous studies [30, 31](Fig. 8a). PfCDPK1 is localized in the parasite membrane marking the periphery of the cytoplasm [30–32]. However, phosphorylated PfCDPK1 is specifically localized towards the apical end of merozoite [33]. PfRKIP is concentrated at the apical end of the merozoites that is exhibited as distinct punctate staining in the developing merozoites within the segmented schizont (Fig. 1c and Fig. 8). The PfRKIP was found to co-localize with PfCDPK1 suggesting that it is present at the apical end of merozoites where PfCDPK1 is also present in its phosphorylated form [33](Fig. 8a and 8b).

**Figure 8.**
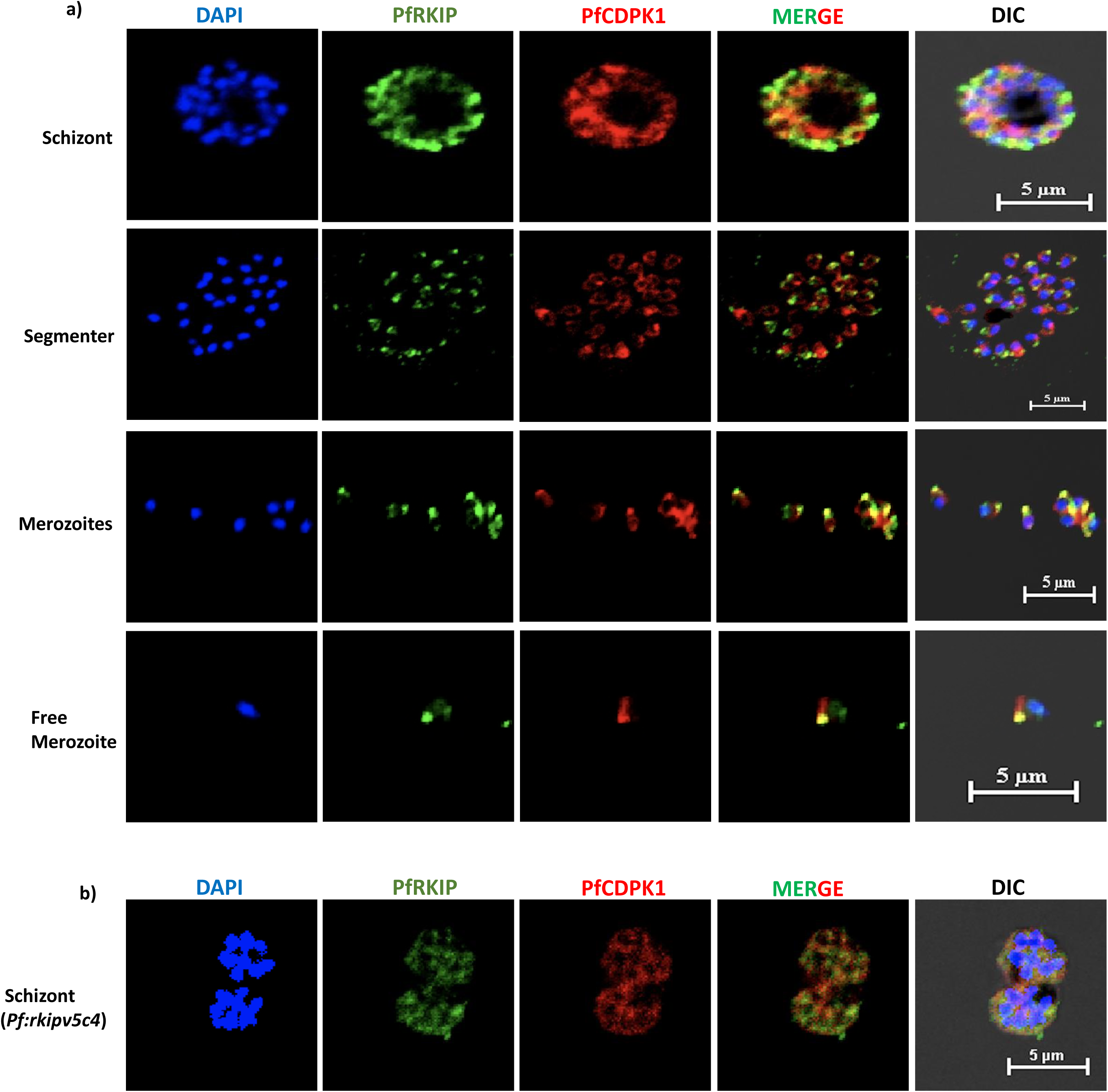
PfRKIP colocalizes with PfCDPK1 within mature schizont and free merozoite. Mature schizonts from WT **(a)** and *pf:rkipv5c4* parasites **(b)** were stained simultaneously with anti-PfRKIP antibody and anti-PfCDPK1 antibody. PfRKIP (green) colocalizes with PfCDPK1 (red) in both the parasites. MERGE image of PfRKIP and PfCDPK1 is shown. The samples were counterstained with 4′,6-diamidino-2-phenylindole (DAPI), a nuclear stain. DIC-Differential Interference Contrast. White scale bar is 5 μm.

To further test the interaction of PfRKIP and PfCDPK1 within the parasite, we took advantage of a proximity biotinylation strategy (Fig. 9b). A modified and improved promiscuous biotin ligase, BioID2 [34, 35] was used to study the interaction between PfRKIP and PfCDPK1. The activity of BioID2 was first verified in *E*. *coli* by expressing the chimeric PfRKIP-BioID2-His as a recombinant protein. The purified PfRKIP-BioID2-His protein resulted in substantial increase in the biotinylation of bovine serum albumin along with itself in the presence of exogenous biotin compared to without biotin condition (Fig. S7a). Purified PfRKIP-His did not show any biotinylation activity either in presence or absence of biotin (Fig. S7a). Also, BSA by itself did not show any biotinylation in presence of biotin (Fig. S7a). These results confirm that BioID2 is functionally active biotin ligase. Therefore, we generated a transgenic parasite that express PfRKIP along with BioID2. BioID2 gene sequence was appended at the 3’ end of *pfrkip* gene in an episomal plasmid along with a V5 epitope tag. The chimeric protein, PfRKIP-BioID2-V5 will biotinylate all the potential proximal and interacting partners within the parasite that can be immunoprecipitated using streptavidin magnetic beads. As a control, we generated a transgenic parasite that overexpressed PfRKIP-cMyc protein from the same plasmid but without BioID2. The presence of the plasmids in the transgenic parasites were verified through a diagnostic PCR (Fig. S7b). Amplification of the target genomic DNA with primer pair, pDC2A/pDC2B (specific for the plasmid DNA flanking the *pfrkip* gene fragment) in the *pfrkip-bioid2-v5* and control (*pfrkip-cmyc*) parasites resulted in amplicons of desired sizes of 1525 bp and 832 bp, respectively (Fig. S7b). The presence of PfRKIP-BIOID2-V5 and PfRKIP-cMyc proteins were further tested in the lysates prepared from *pfrkip-bioid2-v5* and control (*pfrkip-cmyc*) parasites through Western blot. Bands corresponding to the sizes of PfRKIP-BIOID2-V5 and PfRKIP-cMyc were specifically detected by anti-V5 and anti-cMyc antibody in the lysates of *pfrkip-bioid2-v5* and *pfrkip-cmyc* parasites (Fig. 9a). Two parallel bots were probed with anti-β-actin and anti-PfRKIP Ab (Fig. 9a). With anti-PfRKIP Ab, PfRKIP-BIOID2-V5 was detected at higher position (~ 49 kDa) compared to PfRKIP-cMyc (~ 23 kDa) due to the presence of BIOID2 in PfRKIP-BIOID2-V5 (Fig. 9a). Total biotinylated proteins were immunoprecipitated from *pfrkip-bioid2-v5* and control parasites and separated on SDS-PAGE. Western blot of the input sample prepared from *pfrkip-bioid2-v5* and control parasite lysates showed the presence of PfRKIP-BIOID2-V5 and PfRKIP-cMyc with anti-PfRKIP Ab (Fig. 9c, input). With the anti-PfRKIP and anti-PfCDPK1 antibodies, bands corresponding to PfRKIP-BIOID2-V5 (~ 49 kDa) and PfCDPK1 (~ 60 kDa) were detected only in the immunoprecipitated samples in the *pfrkip-bioid2-v5* parasites while no bands at the corresponding positions were detected in the control (Fig. 9c). These results strongly suggest that PfRKIP interacts with PfCDPK1 within the parasites.

**Figure 9.**
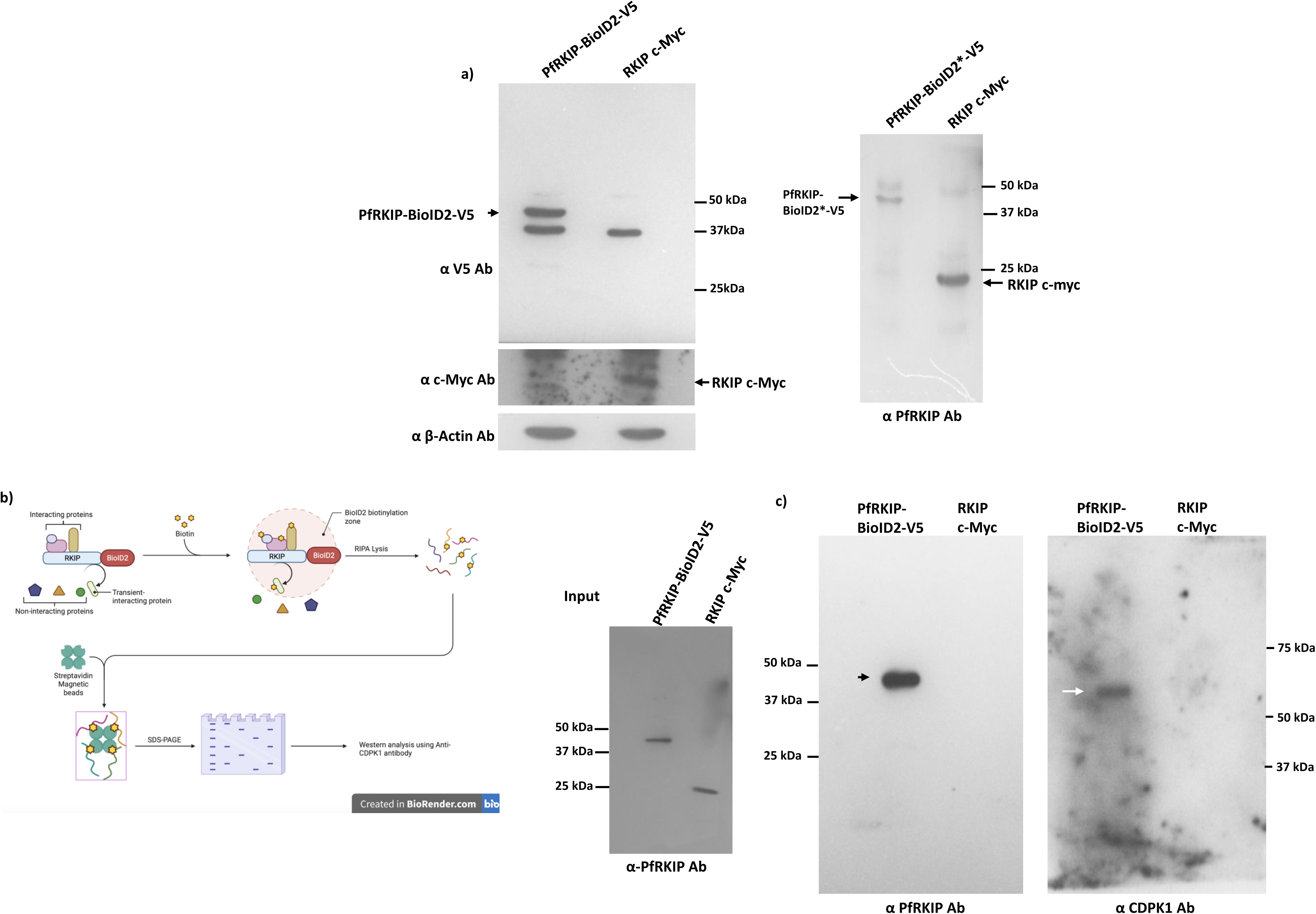
PfRKIP and PfCDPK1 interact with each other within the parasite. We used proximity biotinylation approach to test the interaction of PfRKIP and PfCDPK1 within late stages of the parasite. For this, transgenic parasite with overexpressing PfRKIP-BioID2-V5 were generated along with control parasites overexpressing PfRKIP-cMyc without BioID2. **a)** Parasite lysates prepared from PfRKIP-BioID2-V5 and PfRKIP-cMyc were probed through Western blot with anti-V5 antibody and anti-cMyc antibody. Two parallel blots were probed with anti-beta-actin (α β-Actin Ab) and anti-PfRKIP antibody. **b)** Schematic of the proximity biotinylation approach. The modified bacterial biotin ligase BioID2 (red) is fused to the C-terminus of PfRKIP (with a V5 tag downstream, not shown). BioID2 converts exogenously added free biotin to highly reactive biotinyl-5-AMP which reacts with the primary amines of interacting and proximal proteins leading to their biotinylation (Biotinylation zone). Following biotinylation, the biotinylated proteins were affinity purified from parasite lysates using streptavidin magnetic beads and identified by Western blot using corresponding antibodies. **c)** PfRKIP interacts with PfCDPK1 within the parasite. Western blot showing the detection of ~ 49 kDa PfRKIP-BioID2-V5 and ~ 23 kDa PfRKIP-cMyc (input). Band corresponding to PfRKIP-BioID2-V5 (black arrow) was detected in the immunoprecipitated sample from the *pf:rkipbioid2v5* parasites and not in the control. Band corresponding to the size of PfCDPK1 was detected in the immunoprecipitated sample from the *pf:rkipbioid2v5* parasites with anti-PfCDPK1 antibody (white arrow).

### Pharmacological inhibition of PfRKIP leads to parasite killing

Our results suggest that PfRKIP interacts and colocalizes with PfCDPK1 in mature schizonts and free merozoites. Since PfCDPK1 is important for invasion of red blood cells therefore, we tested whether inhibition of PfRKIP would affect the invasion process. To this end, we used a specific inhibitor of mammalian RKIP called locostatin that blocks key physiological processes in mammalian cells [36, 37]. Before testing the effect of locostatin on the parasite growth, we tested the effect of locostatin on the interaction of PfRKIP and PfCDPK1 through ELISA. PfRKIP was incubated with different concentrations of locostatin and allowed to interact with PfCDPK1 coated on a 96-well plate. The bound PfRKIP was measured using anti-PfRKIP antibodies. The O.D. at 492 nm were plotted against different concentrations of locostatin (Fig. 10a). Interestingly, PfRKIP showed increased interaction with PfCDPK1 when incubated with locostatin compared to without locostatin treatment at all the concentrations of locostatin (Fig. 10a). The increase in the interaction of PfRKIP and PfCDPK1 in presence of locostatin was statistically significant (n=3; p=0.0036, paired t-test) (Fig. 10a). Interestingly, when PfRKIP was not pre-incubated with locostatin, rather it was added concomitantly with other reagents, no effect was observed in the interaction between PfRKIP and PfCDPK1 (Fig. S6) suggesting that pre-incubation of PfRKIP with locostatin causes modification of PfRKIP that allows its tight interaction with PfCDPK1 [25, 38]. Furthermore, locostatin treated PfRKIP showed less interaction with PfCDPK1 in presence of ATP compared to without ATP condition (p= 0.0040, paired t-test; n=3) suggesting that phosphorylation of locostatin-treated PfRKIP reduces its interaction with PfCDPK1 similar to PfRKIP that is not treated with locostatin (Fig. S5 and Fig. 7d). We next tested the effect of locostatin on the growth of parasite under *in vitro* culture conditions. For this, mature schizonts (38-42 HPI) were incubated with increasing concentration of locostatin and ring parasites were counted through Giemsa in the next cycle. With increase in the concentration of locostatin there was a dose dependent decrease in the formation of newly invaded rings (Fig. 10bi) (n=3; p<0.05, paired t-test). Since the defect in new ring formation could be due to block in egress and/or invasion, we counted the number of unruptured schizonts in each treatment condition. The number of unruptured schizonts were comparable to the control till 25 μM locostatin, however at higher concentrations of locostatin there was an increasing trend in the number of unruptured schizonts compared to the control that was not statistically significant (Fig. 10bii). Therefore, the decrease in ring parasitemia in the presence of locostatin may be due to the block in invasion of RBC.

**Figure 10.**
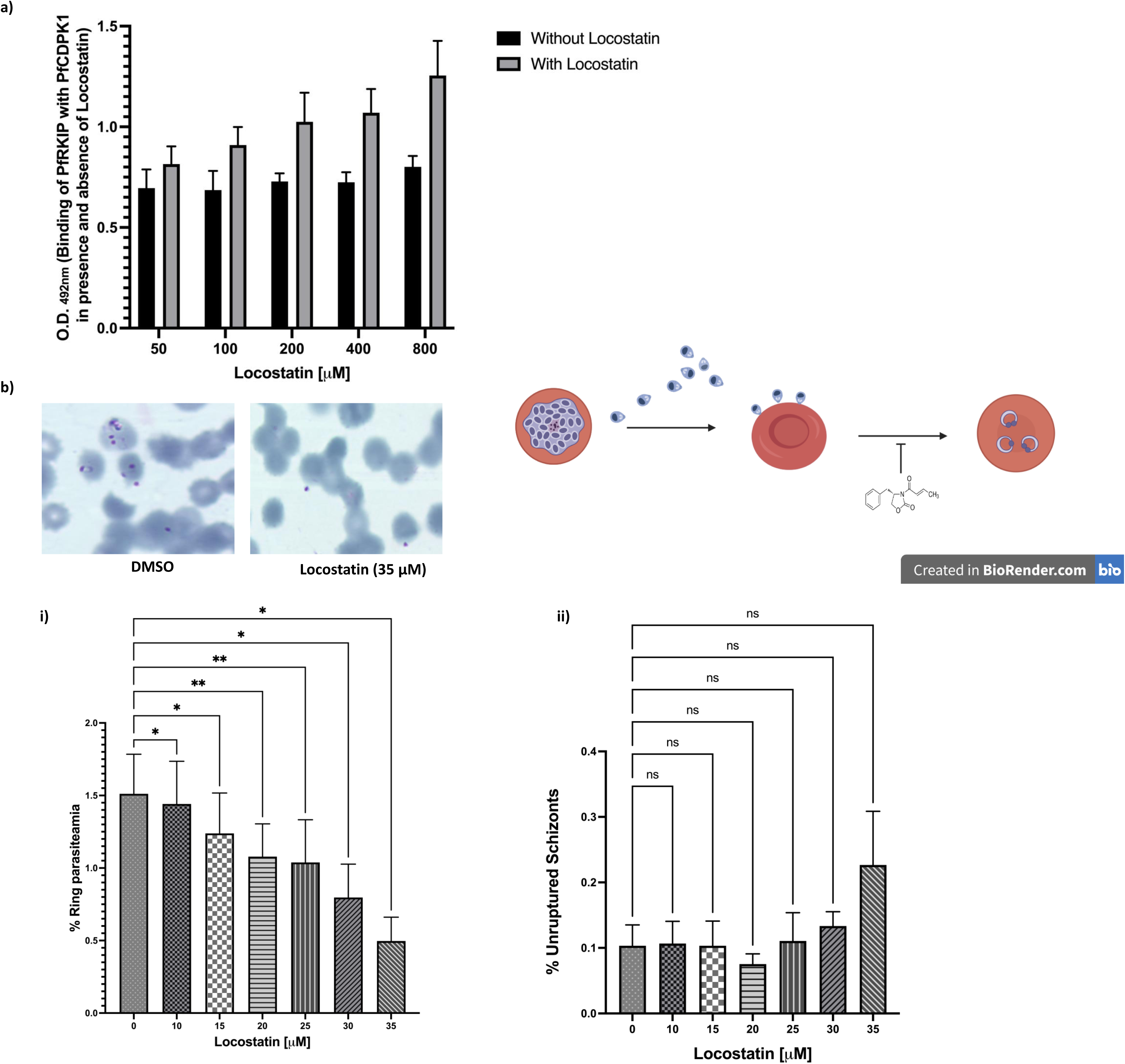
Locostatin increases the interaction of PfRKIP with PfCDPK1 and leads to parasite killing. **a)** Locostatin mediated modification of PfRKIP increases its interaction with PfCDPK1. PfRKIP was treated with different concentrations of locostatin and used to test its interaction with PfCDPK1. With increase in the concentration of locostatin the binding of PfRKIP with PfCDPK1 increased compared to without locostatin. The binding of PfRKIP with and without locostatin on the Y-axis is plotted against the concentration of locostatin (in μM) on the X-axis. The data is obtained from three biological experiments performed in duplicate (n=3, p=0.0036, paired t-test). Error bars represent standard error of mean. **b)** To test the effect of locostatin on parasite egress/invasion of red blood cells, highly synchronized schizonts (38 - 42 HPI) were treated with different concentrations of locostatin and ring parasites along with unruptured schizonts were counted after 12-14 h through Giemsa staining. The number of ring parasites decreased with increase in the concentration of locostatin **(i)**. The % ring parasitemia on the Y-axis is plotted against the locostatin concentration (in μM) on the X-axis **(i)**; **(ii)** the unruptured schizonts on the Y-axis were plotted against the concentration of locostatin (in μM) on the X-axis. The data from three independent biological experiments counted in duplicate was used to plot the graphs in GraphPad Prism 9 (n=3; * and ** denote p<0.05 and p<0.005, respectively; one-way ANOVA). The error bars represent standard error of mean. ns: not significant.

**Figure 11.**
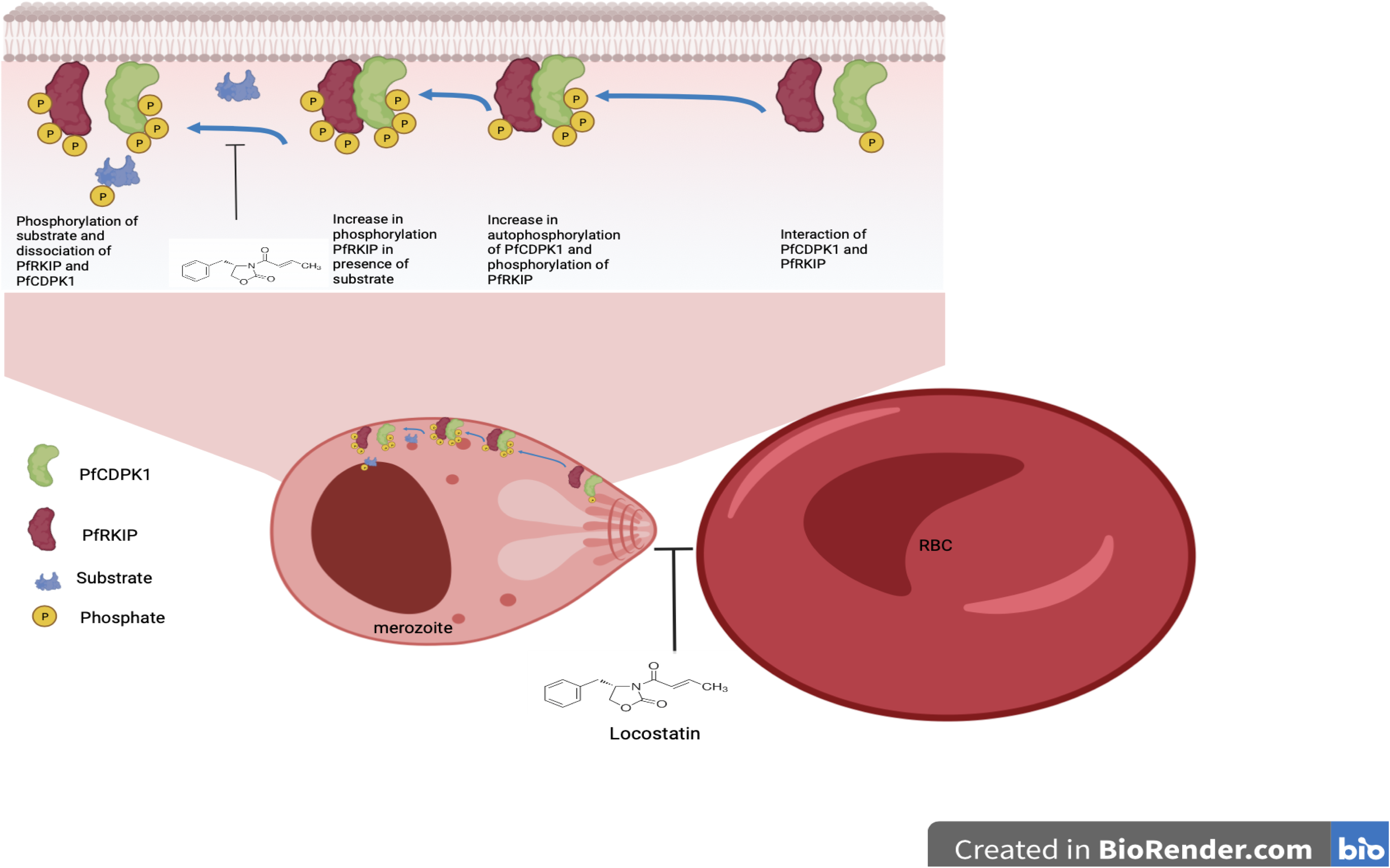
Schematic model for the molecular mechanism of PfRKIP mediated control of PfCDPK1 activity and role of locostatin. Interaction of PfRKIP with PfCDPK1 causes increase in the autophosphorylation of PfCDPK1 that also leads to the phosphorylation of PfRKIP. Phosphorylation of PfRKIP reduces its interaction with PfCDPK1. In presence of a substrate there is further increase in the phosphorylation of PfRKIP that further reduces its interaction with PfRKIP leading to its detachment from PfCDPK1. This allows transphosphorylation of the substrate by PfCDPK1. Locostatin, a pharmacological inhibitor of mammalian RKIP, causes alkylation of PfRKIP that leads to increase in the interaction of modified PfRKIP with PfCDPK1 thereby stabilizing the heterodimeric complex of the two proteins leading to the blockade of PfCDPK1 function. Locostatin mediated inhibition of PfCDPK1 through PfRKIP is perhaps responsible for block in invasion of red blood cells by the malaria parasite.

## DISCUSSION

Raf kinase inhibitor protein in higher eukaryotes is involved in regulation of critical signalling pathways and known to modulate various protein-protein interactions [39–42]. The function of human ortholog of RKIP in *P*. *falciparum* (PF3D7_1219700) has not been explored. Here we have demonstrated that PfRKIP binds with physiologically relevant lipids in a pH dependent manner and importantly, it interacts and regulates the kinase activity of PfCDPK1, a plant like kinase that is important for invasion of red blood cells by *Plasmodium* merozoite.

Our findings reveal pH-dependent binding between PfRKIP and various lipids. Notably, the binding characteristics of PfRKIP are noticeably different from those of HsRKIP. In a significant advancement, we have successfully established the interaction between RKIP and lipids, including phosphatidylethanolamine, using a far-Western technique. Interestingly, the binding between HsRKIP and lipids exhibited a decreasing trend with rise in pH, and intriguingly, no binding was detected at the physiological pH. Conversely, the binding of PfRKIP with PtdIns(3)P and PtdIns(5)P increased with increase in pH. The pH of the buffer containing RKIP increases the flexibility of regions surrounding the ligand binding site that may alter lipid specificity [18]. Moreover, histidine residues in and around the pocket are sensitive to pH induced conformational changes that may further contribute towards specificity in lipid binding and protein-protein interactions [18]. Mammalian RKIP has been crystallized at low pH with high concentration of phosphorylethanolamine (the head group of phosphatidylethanolamine) however, at near physiological conditions there is no report of its binding with the lipid moieties. The lipid binding site in RKIP is present on the surface of the protein and is constituted by a small region at the C-terminal helix and two regions defined as CR1 and CR2. The CR1 and CR2 consist of a few conserved residues while the C-terminal helix is highly variable among different PEBP proteins. Variant residues in CR1 and CR2 dictate the size and shape of the ligand binding site that perhaps is an important determinant of specificity of PEBP proteins for different ligand. Furthermore, a strip of basic residues surrounding the ligand binding site helps in stabilization of the anionic group of the lipid moiety. In PfRKIP, some of the basic residues in the strip are conserved while few positions are constituted with acidic residues that may further contribute towards distinct lipid interaction profiles. One of the two cis-peptide bonds present in mammalian PEBP proteins is also conserved in PfRKIP (K88-E89) and is postulated to be important for membrane interactions. PfCDPK1 is critical for the formation of both male and female gametes. Disruption of PfCDPK1 in a mutant parasite background show upregulation of transcripts for the genes associated with the sexual stage development [43]. In addition, the transcript of PfRKIP is also upregulated (~2.5 fold) in the PfCDPK1 KO parasite. Interestingly, overexpression of a putative PEBP-RKIP family member in plants show early flowering phenotype compared to the wild type and also led to the upregulation of apetala transcription factor [44]. Increase in the transcripts of sexual stage genes in PfCDPK1 KO parasites maybe an indirect consequence of PfRKIP upregulation or directly related to de-repression of sexual stage genes due to PfCDPK1 KO.

Mutation of conserved amino acids such as P80 and H92 (part of CR1) to Leu and Ala, respectively in PfRKIP results in decrease in interaction of PfRKIP with PtdIns(3)P suggesting that the conserved amino acids in the conventional lipid binding domain are important determinants for binding with lipids. Our bioinformatics analysis showed the presence of P80 near the residues that interact with PtdIns(3)P suggesting that although P80 was not directly involved in non-covalent interactions with PtdIns(3)P it may still have a stabilizing effect on the lipid and protein interaction. As such we found maximum reduction (10 fold) in lipid binding with the P80L mutant. The ITC experiments for the study of thermodynamic parameter of lipid and RKIP interaction revealed biphasic and monophasic interaction of PfRKIP and HsRKIP, respectively with PtdIns(3)P. This suggest that the Plasmodium RKIP may contain two different lipid binding sites. Our bioinformatics study also suggests that a lipid binding site is present towards the C-terminus of PfRKIP in addition to the conventional site present in HsRKIP. *Plasmodium* RKIPs have a unique representation of a left-handed alpha helix (α5) towards the C-terminus that is not present in HsRKIP. Deletion of α5 in PfRKIP led to decrease in interaction with PtdIns(3)P showing importance of the left-handed helix in lipid binding. Left handed alpha helix is not a common structural feature and is found in only few structures deposited in PDB. The left-handed α helix maybe required for binding with the lipid head groups that are exposed towards the cytosolic phase. It is interesting to note that PfRKIP^Δα5^ mutant did not show any effect in interaction with PfCDPK1 suggesting that α5 is perhaps involved in recognition of lipids moieties and is not involved in the interaction of RKIP with its partners proteins. It will be interesting to further explore the significance of having two lipid binding domains in PfRKIP as opposed to one in HsRKIP.

RKIP binds to Raf1 and inhibits Raf1/MEK/ERK signalling pathway. Upon phosphorylation at S153 by PKC, RKIP dissociates from Raf1 and activates Raf1/MEK/ERK signaling pathway and concomitantly results in protein kinase A activation through inhibition of G-protein coupled receptor kinase 2 [10, 40, 45]. Interestingly, PKA mediated phosphorylation of RKIP increases its propensity to get phosphorylated at S 153 by PKC thus establishing a positive feedback-loop [45]. Our results show that PfRKIP binds with PfCDPK1 and increases the autophosphorylation of PfCDPK1. The autophosphorylation of PfCDPK1 could be the result of a positive feedback loop established through PfCDPK1 mediated phosphorylation of PfRKIP. Autophosphorylation of CDPKs is known to increase the kinase activity of the protein that allows phosphorylation of downstream substrates by CDPKs [46]. In addition to being directly required for substrate phosphorylation, autophosphorylation of CDPK is also an important determinant for mediating interaction between different proteins in signalling cascades [47]. In this role CDPK performs dual function of a scaffold and a kinase that allows binding of two proteins out of which phosphorylation of one is required for the formation of a heterodimeric complex [47]. Likewise, CDPKs may also mediate formation of multimeric complexes. Interestingly, PfCDPK1 has been shown to be a part of high molecular weight complex in schizonts and merozoites [33, 48]. PfCDPK1 is localized on the inner leaflet of parasite membrane and other membranous structures and is also associated with the motor complex where it helps in the phosphorylation of inner membrane complex and glideosome associated proteins [31, 32, 49]. Interestingly, however phosphorylated CDPK1 has been shown to get preferentially localized at the apical end of merozoites within late schizonts and free merozoite however, not within any of the apical organelles. The strategic localization and phosphorylation of IMC and glideosome associated proteins coincides very well with the function of PfCDPK1 during invasion of RBCs by merozoites [43, 48, 49]. As such disruption of PfCDPK1 by conditional knock-down or complete knock-out in mutant parasite background has been shown to affect invasion of RBCs [43, 49]. Interestingly, PfRKIP shows punctate staining in late schizonts that co-localizes with PfCDPK1 perhaps with the phosphorylated form of PfCDPK1 at the periphery. We did not have access to the phospho-specific anti-PfCDPK1 antibodies therefore, we could not perform co-localization study of phosphorylated PfCDPK1 and PfRKIP. Nevertheless, coexistence of PfRKIP and PfCDPK1 at the apical end of the merozoite might be required for the activation of PfCDPK1 that is important for invasion of red blood cells.

There is no potential phosphorylation site in PfRKIP corresponding to S153 in HsRKIP. However, S96 in PfRKIP was found to be phosphorylated by recombinant PfCDPK1 and was therefore proposed as a potential putative phosphorylation site within the parasite. We have found that blocking the phosphorylation of S96 residue in PfRKIP by substitution with ALA (S96A) decreases the overall phosphorylation of the mutant PfRKIP by PfCDPK1 compared to the WT PfRKIP suggesting that S96 is phosphorylated in PfRKIP however, there are other potential sites that are phosphorylated by PfCDPK1 under *in vitro* conditions. We have not explored the effect of blocking the phosphorylation of S96 on the parasite growth moreover, an authentic phosphorylation site in PfRKIP within the parasite needs to be investigated. Leukotriene D4 treatment of epithelial cells results in protein kinase C (PKC) mediated phosphorylation of RKIP that causes activation of PKA signalling cascade [50]. In the malaria parasite, PKA mediated phosphorylation of cytoplasmic tail of an invasion related gene, apical membrane antigen 1 (AMA1) is critical for invasion of RBCs [51, 52]. It might be possible that RKIP regulates other unexplored signalling pathways in the malaria parasite.

Locostatin, a specific pharmacological inhibitor of mammalian RKIP, occupies the ligand binding site of PEBP proteins and causes alkylation of a histidine residue in the PEBP domain. Locostatin inhibits the ability of RKIP to interact with Raf1 and as such the activity of Raf1 is not blocked by RKIP in presence of locostatin leading to the activation of MAPK pathway [37, 38]. However, locostatin modified RKIP do not show appreciable disruption with IKKα, another RKIP binding protein [25](beshir). Surprisingly, our data shows that locostatin mediated modification of RKIP causes increase in the interaction between PfRKIP and PfCDPK1. In the *in silico* data on interaction of PtdIns(3)P with PfRKIP, we did not found H92 residue in the vicinity of the lipid however, HIS at the corresponding position (H86) in HsRKIP was found to interact with the lipid. Therefore, it could be possible that although H92 is involved in lipid binding, it may be majorly involved in interaction of PfRKIP with other proteins. Hence neutralization of H92 through locostatin mediated alkylation abrogates unfavourable ionic interaction between PfRKIP and PfCDPK1 leading to increased interaction between the two proteins. Interestingly, the increase in the interaction between the two proteins was obtained only when RKIP was pre-incubated with locostatin. When added along with the two proteins, locostatin did show any effect on the interaction of RKIP and CDPK1. This would mean that the locostatin binding site was masked during the interaction of CDPK1 and RKIP and it was not able to alkylate the histidine residue. H92 is involved in interaction between RKIP and CDPK1 is supported by reduced interaction of H92A RKIP with CDPK1 in the ELISA based assay. Due to locostatin mediated increase in the interaction between the 2 proteins, PfCDPK1 may not be able to phosphorylate its downstream substrates. Parasites treated with increasing concentration of locostatin showed dose dependent decrease in ring parasitemia that could perhaps be due to the formation of a tight heterodimeric complex between PfRKIP and PfCDPK1 leading to sequestration of PfCDPK1. Interestingly, the percentage of unruptured schizonts were similar in the presence and absence of locostatin suggesting that inhibition in ring formation due to locostatin treatment was due to the block in invasion and not egress of merozoites. Specific compounds may be designed based on locostatin that increase the interaction of PfRKIP with PfCDPK1. These compounds can be used in combination with existing anti-malarial drugs for better control of malaria infection.

## EXPERIMENTAL PROCEDURES

### Cloning of full-length PfRKIP and HsRKIP

The full-length *rkip* (573bp; PlasmoDB ID: PF3D7_1219700) was amplified using cDNA prepared from asynchronous *P*. *falciparum* asexual blood stage parasites with the primer set: PfRKIPet42_F and PfRKIPet42_R (see Table 3 for primer sequence). The full-length *rkip* gene from humans (573bp) was amplified using cDNA prepared from HEK-293 cells with the primer set: HsRKIPet42_F2 and HsRKIPet42_R. High-fidelity PrimeStar HS DNA polymerase was used to set up the amplification reaction and the following PCR conditions were employed: 98 °C/2 min, [98 °C/10 sec, 52 °C/20 sec, 62 °C/30 sec] x36, 62 °C/10 min. Amplified DNA fragment was cloned using conventional molecular cloning procedure between NdeI & XhoI sites in pET42a(+) plasmid. The recombinant DNA plasmids were then transformed into *E*. *coli* DH5-α competent cells. Positive colonies containing the desired GoI were confirmed by PCR, double restriction digestion of the recombinant plasmids and DNA sequencing.

**Table 3.**
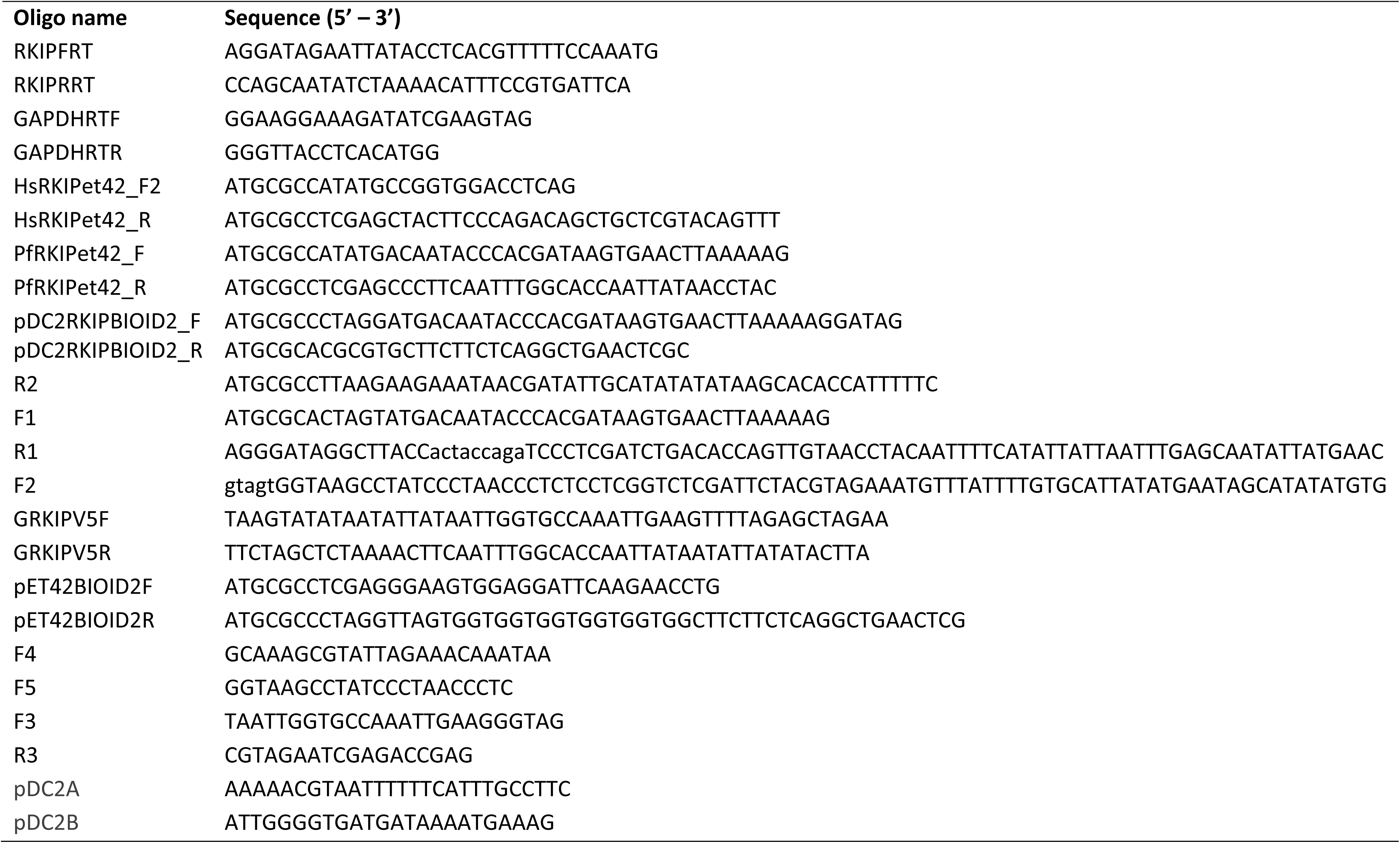
DNA sequence (5’-3’) of the oligos used in the study.

### Expression and Purification of recombinant proteins

The sequence verified plasmid construct containing *pfrkip* in pET42a+ vector were transformed into *E. coli* BL21(DE3)pLysS competent cells for protein expression. PfRKIP was expressed as a chimeric protein with a 8X His-tag at C-terminus. Protein expression was induced with 1 mM IPTG for 10 h at 24 ^0^C. The *E*. *coli* cells were pelleted at 5000 g for 10 min at 4 ^0^C. The pelleted cells were lysed in a lysis buffer with the composition: 1 mM PMSF, 10 mM Tris-HCl, pH 7.5, 100 mM NaCl, 0.1 mg/mL lysozyme & 10 mM MgCl_2_, 10 % glycerol followed by sonication for 10 min (15 sec pulse ON, 20 sec pulse OFF, 10 % amplitude). The lysate was then centrifuged at 12,000 g for 1 h. The clear lysate containing recombinant PfRKIP was incubated with Ni-NTA sepharose beads O/N at 4 ^0^C. The beads were allowed to settle by centrifuging at 500 g for 2 min at 4 ^0^C followed by extensive washing with the wash buffer with the following composition:10 mM Tris-Cl, pH 7.5 & 150 mM NaCl containing either 20- or 50 mM imidazole. The chimeric PfRKIP-HIS was eluted from the Ni-NTA sepharose beads with different concentrations of imidazole in the wash buffer. The fractions were run on SDS-PAGE. Fractions with similar purification profile were poled together followed by buffer exchange with 50 mM Tris-Cl, pH 7.5 & 150 mM NaCl and stored at −80 ^0^C for various experiments. The identity of the recombinant PfRKIP was verified by mass spectrometry.

### Expression and Purification of HsRKIP

The sequence verified plasmid construct containing *hsrkip* gene in pET42a+ vector was successfully transformed in *E*. *coli* BL21(DE3)pLysS competent cells for protein expression. HsRKIP was expressed as a chimeric protein with His-tag at C-terminus. *E*. *coli* cultures were grown in LB media with appropriate antibiotics at 37 °C. Recombinant HsRKIP protein expression was induced with 1 mM IPTG at an OD_600_ of 0.8. The recombinant protein expression was continued for 5 h at 30 °C. The bacterial cells were harvested at 5000 g by centrifugation and resuspended in lysis buffer of the following composition: 1 mM PMSF, 10 mM Tris-HCl, pH 7.5, 100 mM NaCl, 0.1 mg/mL lysozyme & 10 mM MgCl_2_, 10 % glycerol. The bacterial cells were allowed to be incubated in the lysis buffer for 1 h on ice followed by sonication for 10 min (pulse: 15 sec ON, 20 sec OFF, 10 % amplitude). The lysate was then centrifuged at 12,000 g for 1 h, and the clear supernatant with recombinant HsRKIP was incubated at 4 °C with Nickel-NTA Sepharose beads (GE-healthcare) for 4 h. The Ni-NTA resins with bound HsRKIP were successively washed with 50 mM Tris-Cl, pH 7.5 & 300 mM NaCl containing either 20 or 50 mM imidazole. The bound HsRKIP recombinant protein was eluted in elution buffer (50 mM Tris-Cl, pH 7.5 & 300 mM NaCl) containing 100 mM, 200 mM, 250 mM and 500 mM imidazole. Proteins with similar SDS-PAGE profile were poled together followed by buffer exchange with 50 mM Tris-Cl, pH 8.0 & 150 mM NaCl and stored at −80 ^0^C for various experiments. The recombinant HsRKIP protein was verified by Western blot using anti-His antibodies and also confirmed by mass spectrometry.

### Cloning and expression of recombinant PfRKIP mutant proteins

Five different mutants of PfRKIP were designed based on previous literature and alignment with HsRKIP and PvRKIP protein sequence. The following mutations were designed: P80L, H92A, S96A, K143A, and Δα5 (for primer DNA sequences that were used for site-directed mutagenesis see Table 4). The mutations were incorporated in WT *pfrkip* DNA sequence using QuikChange XL site-directed mutagenesis kit (Stratagene, USA) following the manufacturer’s instructions. Primer pairs containing the designed modification were designed using Agilent QuickChange Primer Design software [https://www.agilent.com/store/primerDesignProgram.jsp]. Recombinant plasmids containing the desired mutation were transformed into *E. coli* DH5α and the mutations were confirmed by DNA sequencing.

**Table 4.**
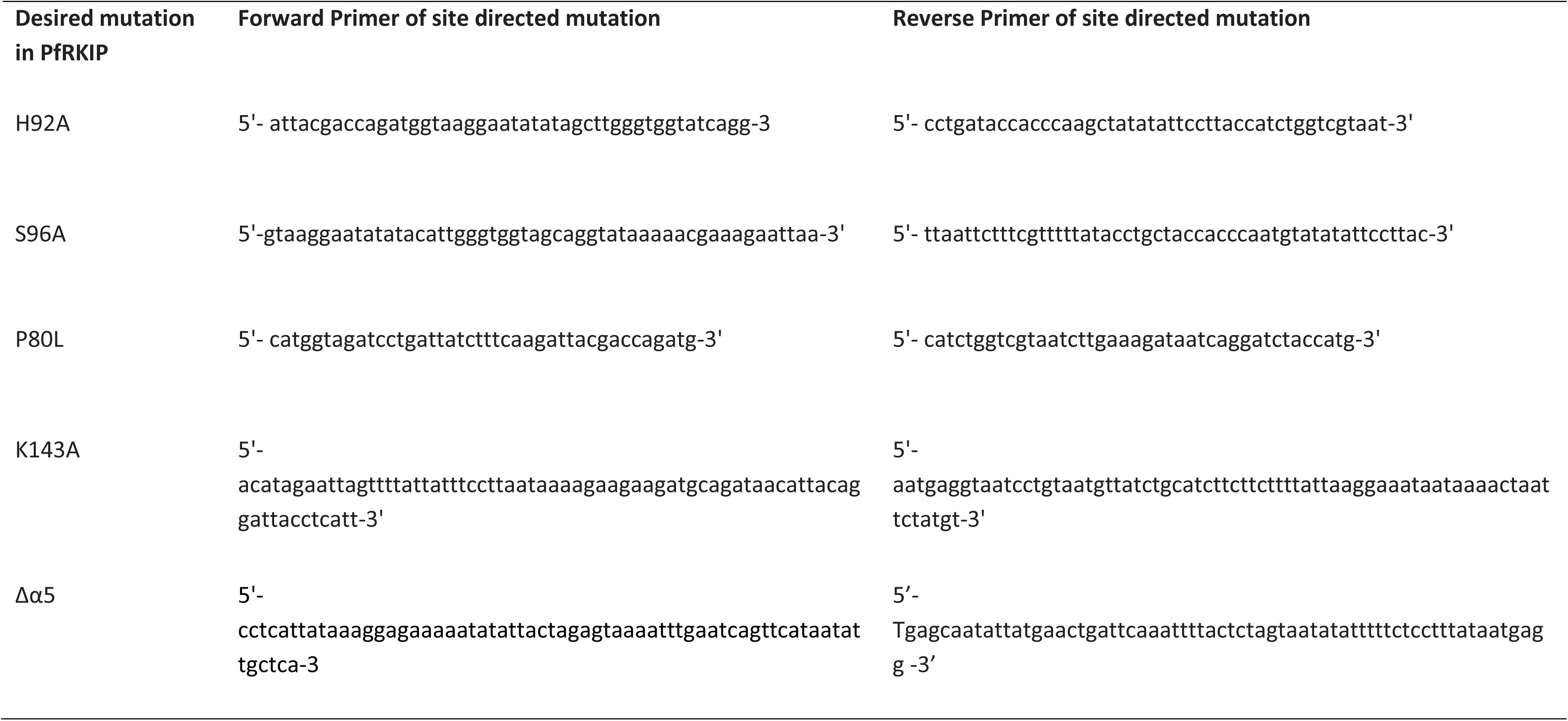
Primer used for site directed mutagenesis. Five mutations were introduced in the PfRKIP protein sequence: P80L, H92A, S96A, K143A, and Δα5 using the listed primer pairs. The primer pairs were designed using the Agilent QuickChange Primer Design software.

**Table 5.** Thermodynamic parameter for the binding of proteins with lipid from ITC experiment.

| SYSTEM | K<sub>1</sub><br>(M<sup>-1</sup>) | ΔH<sub>1</sub><br>(kcal/mol) | TΔS<sub>1</sub><br>(kcal/mol) | ΔG<sub>1</sub><br>(kcal/mol) | K<sub>2</sub><br>(M<sup>-1</sup>) | ΔH<sub>2</sub><br>(kcal/mol) | TΔS<sub>2</sub><br>(kcal/mol) | ΔG<sub>2</sub><br>(kcal/mol) |
| --- | --- | --- | --- | --- | --- | --- | --- | --- |
| PfRKIP | 8.7 × 10<sup>4</sup><br>(±2.4 × 10<sup>4</sup>) | -19.9 (±1.5) | -13.2 | -6.7 | 4.4 × 10<sup>4</sup><br>(±8.9 × 10<sup>3</sup>) | -49.7 (±6.2) | -43.5 | -6.5 |
| PfRKIP Δα5 | 5.6 × 10<sup>4</sup><br>(±1.8 × 10<sup>4</sup>) | -28.4 (±3.6) | -21.9 | -6.5 | 4.9 × 10<sup>4</sup><br>(±1.3 × 10<sup>4</sup>) | -49.2 (±8.9) | -42.6 | -6.6 |
| PfRKIP H92A | 6.4 × 10<sup>4</sup><br>(±1.3 × 10<sup>4</sup>) | -40.0 (±3.0) | -33.4 | -6.6 | 4.2 × 10<sup>4</sup><br>(±1.1 × 10<sup>4</sup>) | -64.4 (±12.0) | -58.1 | -6.3 |
| HsRKIP | 2.3 × 10<sup>4</sup><br>(±1.9 × 10<sup>4</sup>) | -11.7 (±6.6) | -5.78 | -5.9 | 5.5 × 10<sup>3</sup><br>(±4.1 × 10<sup>3</sup>) | -3.5 (±18.8) | 1.60 | -5.1 |
| PfRKIP S96A | 6.4 × 10<sup>4</sup><br>(±8.3 × 10<sup>3</sup>) | -15.5 (±0.8) | -8.97 | -6.5 | 4.7 × 10<sup>4</sup><br>(±1.1 × 10<sup>4</sup>) | -21.1 (±3.3) | -14.8 | -6.4 |
| PfRKIP P80L | 9.2 × 10<sup>3</sup><br>(±1.5 × 10<sup>3</sup>) | -89.6 (±11.7) | -84.1 | -5.5 | 3.3 × 10<sup>3</sup><br>(±6.0 × 10<sup>2</sup>) | -99.9 (±44.6) | -95.10 | -4.8 |

The sequence verified pET42a(+) plasmid constructs containing the desired mutation in *pfrkip* were further transformed in *E. coli* ArcticExpress(DE3)RIL competent cells for protein expression. The mutant PfRKIP proteins were expressed as chimeric protein with 8xHis tag at C-terminus and induced for protein expression with 1 mM IPTG for 24 h at 12 ^0^C. The recombinant PfRKIP mutant proteins were purified as described for the WT PfRKIP.

### Generation of antisera against recombinant PfRKIP

For generation of antisera against recombinant PfRKIP, we used Sprague Dawley rats. The study was duly approved by the Institutional Animal Ethics Committee (IAEC code # 20/2019) and the study was conducted through Central Laboratory Animal Resources (CLAR), Jawaharlal Nehru University. The rats were immunized subcutaneously for generating antisera against recombinant PfRKIP. The following immunization schedule was followed: priming of the animals was done with 100 µg of recombinant PfRKIP protein formulated in equal volume of complete Freund’s adjuvant. First booster immunization formulated with 50 µg recombinant PfRKIP in saline with equal volume of incomplete Freund’s adjuvant was given at after 3 weeks of priming while 2^nd^ booster was given 3 weeks later to the 1^st^ booster dose. The terminal bleed was drawn from the immunized animals after 1 week of the 2^nd^ booster. The antisera along with the pre-bleed were stored in small aliquots for later usage.

### Detection of PfRKIP in the parasite by Western blot

The presence of PfRKIP in asexual blood stages was detected by Western blotting. *P*. *falciparum* blood stage culture was tightly synchronized by treatment of ring stage parasite with two consecutive cycles of 5 % sorbitol. Highly synchronized ring, trophozoite and schizont stage parasites were harvested by centrifugation at 500 g for 5 min at RT. The parasites were treated with 0.05 % saponin in PBS for 10 min in ice followed by centrifugation at 500 g for 5 min at 4 ^0^C. Three washes of 1xPBS were given to remove the lysed RBC components and the saponin. Finally, the parasites devoid of RBC were resuspended in radio-immunoprecipitation assay buffer (50 mM Tris, pH 8.0, 150 mM NaCl, 1 mM EDTA, 1 mM EGTA, 1 % Nonidet P-40, 1 % sodium deoxycholate, protease inhibitor cocktail (Roche Applied Science)), and incubated in ice for 1 h with intermittent shaking. The detergent-resistant and detergent-soluble fractions were separated by centrifugation at 15,000 *g* for 30 min at 4 ^0^C. Both fractions were separated by SDS-PAGE and transferred to PVDF membrane for Western blotting. The PVDF membrane was blocked with 5 % skimmed milk in TBST buffer followed by incubation of the membrane in blocking buffer containing anti-PfRKIP antiserum (1:2000 dilution) for either O/N at 4 ^0^C or 1 h at RT. The membrane was washed 3X with 1x TBST for 5 min each followed by incubation with anti-rat IgG secondary antibody conjugated with horseradish peroxidase (HRP) (Sigma) (1:2000 dilution) for 1 h at RT. The membrane was treated with ECL plus Western blotting detection kit femtoLUCENT^TM^ PLUS HRP GBiosciences and the signal for the presence of PfRKIP was detected by exposing the membrane onto the X-ray for different time period. Anti-β-actin antibodies (Sigma, 1:5000 dilution) were used as a loading control for the Western blot experiments.

### Immunofluorescence Assay (IFA)

Immunofluorescence assay was used to test the localization of PfRKIP in different asexual blood stages of malaria parasite. The protocol used for IFA is as described in a previous study [43]. Briefly, thin smear of different stages of synchronized *P. falciparum* blood stage parasite were made on glass slides, air-dried, and fixed in prechilled methanol for 20 min. The slides were blocked with 5 % BSA in 1xPBS at room temperature for 1 h followed by incubation with anti-PfRKIP rat serum (1:100 dilution) in the blocking buffer. The slides were washed three times with 1xPBS for 5 min each on a rocker. The slides were then incubated with secondary antibody, goat anti-rat IgG Alexa Fluor 488 (1:200 dilution). The slides were washed three times with 1xPBS for 5 min each on a rocker. The slides were then incubated with 1 ug/ml of 4’,6-diamidino-2-phenylindole (DAPI) (Invitrogen) in 1xPBS for 20 min at RT. The slides were finally washed three times with 1xPBS for 5 min each on a rocker. Finally, the slides were mounted with ProLong Gold antifade reagent (Invitrogen, USA) and analyzed using a Nikon A1 confocal microscope. For co-localization experiments slides with smear of *P. falciparum* schizont/merozoites were simultaneously incubated with anti-PfRKIP rat serum (1:100) and anti-PfCDPK1 rabbit serum (1:500 dilution). The secondary antibodies used for the co-localization experiments were goat anti-rat IgG conjugated with Alexa Fluor 488 (1:500) and goat anti-rabbit IgG conjugated with Alexa Fluor 594 (1:800).

### Parasite *in vitro* culture and maintenance

The NF54 strain of *Plasmodium falciparum* was grown at 2 % haematocrit in O+ human RBCs (Rotary Blood Bank, New Delhi, India) in RPMI1640 medium containing L-glutamine and supplemented with 25 mM HEPES (MilliporeSigma, USA), 50 µg/ml Hypoxanthine (MilliporeSigma, USA), 5 % heat inactivated, A+ human sera (Rotary Blood Bank, New Delhi, India), 0.25 % Albumax I (ThermoFisher Scientific, USA), 25 mM sodium bicarbonate (SigmaAldrich, USA) and 10 ug/ml gentamycin (Gibco, ThermoFisher Scientific, USA) as described earlier (46). The parasites were grown in a gaseous environment with the following composition: 5% O_2_, 5% CO_2_, and N_2_ at 37 °C. For collection of highly synchronized asexual blood stages, the parasites were synchronized by 5 % sorbitol followed by percoll:sorbitol enrichment of mature stage parasites. The purified late stage parasites were allowed to invade for 4 h in fresh RBCs followed by sorbitol treatment. Highly synchronized ring (9-13 HPI), trophozoite (32-36 HPI) and schizont (44-48 HPI) stage parasites were collected after 9 h, 32 h, and 44 h for Western blot experiments. For RNA isolation from synchronized parasites, the parasite pellet was treated with 0.01 % saponin in 1xPBS for 10 min in ice followed by 2 washes of 1XPBS for 5 min each. The synchronized parasite pellet devoid of RBC was resuspended in TRIZOL solution (Invitrogen, USA) and kept at −80 ^0^C till further processing.

### Generation of transgenic parasites

We generated a transgenic parasite expressing endogenous PfRKIP protein with a C-terminal V5 epitope tag. The endogenous *pfrkip* locus was targeted by cloning 20 nucleotide guide region (5’-ATAATTGGTGCCAAATTGAA-3’) with primer pair GRKIPV5F/ GRKIPV5R using In-Fusion (Clontech, Mountain View, CA) in pL6eGFP plasmid [53] generating pL6eGFPgV5 plasmid. The homology arm of 1024 nucleotides corresponding to the 3’ DNA sequence of *pfrkip* followed by V5 epitope tag and 3’ UTR region of *pfrkip* was constructed by using two overlapping PCR with the primer sets: F1/R1 and F2/R2. The final homology arm was cloned in pL6eGFPgV5 plasmid between SpeI and AflII making the final construct pL6eGFPgV5RKIPV5 plasmid. Cas9 endonuclease was expressed from the plasmid PUF1 that was co-transfected along with pL6eGFPgV5RKIPV5 for the desired tagging of *pfrkip* with V5 epitope [53, 54]. The PfRKIP::V5 transgenic parasites were cloned by limiting dilution and 4 individual clones were verified by diagnostic PCR. The F4/R2 primer pair was used to PCR amplify the modified *pfrkip* locus and the desired modification was verified through DNA sequencing using internal primers. The transgenic parasite clone C4 was further verified for the expression of V5 tagged PfRKIP through Western blotting with anti-V5 antibodies.

For co-immunoprecipitation of PfRKIP, we utilized a proximity biotinylation strategy. The activity of the modified biotin ligase called BioID2 was first verified by expressing PfRKIP-BioID2 chimeric protein in *E*. *coli*. To this end, the BioID2 DNA sequence was PCR amplified from the plasmid pm2gt-hsp101-bioid2 (kind gift from Josh Beck) using the primer pair: pET42BIOID2F and pET42BIOID2R. The PCR amplified fragment was cloned in frame with *pfrkip* gene in pET42a(+) with 6X-His tag at the C-terminus between XhoI and AvrII restriction enzyme sites. Double restriction digestion positive clones were sequence verified and used for the transformation of BL21(DE3)pLysS competent cells for expression of the chimeric protein. PfRKIP-BioID2 recombinant protein was purified following the methodology as described for the purification of recombinant PfRKIP. Biotinylation experiments with the chimeric recombinant PfRKIP-BioID2 were performed for confirming the activity of the BioID2 as described in the later section. The pfrkip-bioid2 DNA sequence from pET42a(+) was amplified using the primer set: pDC2RKIPBIOID2_F and pDC2RKIPBIOID2_R. The PCR amplified product was cloned between AvrII and MluI restriction enzyme sites in the plasmid, pDC2-CDPK1V5 [43]. The double digestion positive clones were verified through DNA sequencing. One of the sequence verified clones of pDC2RKIP-BioID2V5 was transfected in WT parasite using the conditions as described above. BSD positive parasites containing pDC2RKIP-BioID2V5 plasmid were verified through a diagnostic PCR using plasmid specific primer set, pDC2A and pDC2B. As a control for the biotinylation experiments, we generated a transgenic parasite that overexpressed pfrkip-cMyc without BioID2. Presence of the plasmids pDC2RKIP-BioID2V5 and pDC2RKIP-cMyc in the BSD positive parasites were further confirmed through the detection of PfRKIP protein through Western Blotting with anti-V5 and anti-cMyc antibodies, respectively.

### Lipid Binding Assay with recombinant PfRKIP and HsRKIP

To mimic different pH conditions, the recombinant PfRKIP and HsRKIP proteins were resuspended in appropriate buffers. For pH 4 protein solutions were made by dissolving 20 ug/ml of PfRKIP and HsRKIP proteins in 0.2 M sodium acetate buffer (pH 4). For pH 6.5 and pH 7.5, 20 ug/ml of PfRKIP and HsRKIP proteins were dissolved in 150 mM NaCl and 50 mM Tris-Cl pH6.5 and pH 7.5, respectively.

PIP MicroStrips^TM^ membranes (P23752, ThermoFisher Scientific) were blocked in TBS-T buffer containing 3% fatty acid–free BSA with gentle rocking for 1 h at room temperature. Then the membranes were incubated with 20 ug/ml of PfRKIP and HsRKIP protein solution of pH 4.0, pH 6.5 and pH 7.0 respectively for 4 h at RT. The membrane was washed with TBS-T + 3 % fatty acid–free BSA three times using gentle agitation for 10 min each and incubated with anti-His antibody at a 1:5000 dilution in blocking buffer for overnight at 4 °C followed by 3 washes with TBST without BSA. The membrane was then incubated with anti-Rabbit secondary antibody conjugated with horseradish peroxidase in the blocking buffer for 1 h at room temperature followed by 3 washes with TBST. The blot was developed in chemiDOC XRS+.

### Molecular Docking studies of lipids with PfRKIP

To gain a deeper understanding of the distinct interaction mechanisms involved in PfRKIP/HsRKIP and lipids under varying pH conditions, we adopted Molecular docking studies.

### Protein Preparation

The two proteins, namely PfRKIP (PDB ID-2R77), and HsRKIP (PDB ID-1BD9) were retrieved from Protein Data Bank (PDB). Proteins were subjected to protein preparation before molecular docking using Protein Preparation Wizard in Maestro platform of Schrodinger software. During the pre-process steps of protein preparation, addition of hydrogen bonds, creation of zero-order bonds to metals and disulphide bonds, conversion of selenomethionines to methionines, filling up of missing side chains and missing loops using prime, addition of cap termini and deletion of water molecules beyond 5 Å were carried out. The pre-processed proteins were then taken to H-bonds refinement and Optimization processes. The structures are visualized using [55], the co-crystallized ligands were removed before proceeding with the molecular docking studies. Finally, the proteins were subjected to energy minimization by using the force field OPLS_2005 [56, 57]. The PROPKA value was considered as the value of the pH at which each of those proteins were crystallized.

For PfRKIP, we were interested in comprehending the alterations in molecular interactions between the protein and the chosen ligands at the physiological pH (pH 7.5). To satisfy this criterion, pH as well as PROPKA values were set to 7.5 during the protein preparation.

### Ligand preparation

SDF (Structure-Data File) files of all the ligands including Phosphoryl Ethanolamine (PubChem CID - 129735059), Phosphatidylinositol-3,4,5-trisphosphate (PubChem CID - 53477782), Phosphatidic acid (PubChem CID - 446066), and Phosphate (PubChem CID - 1061) were collected from PubChem database. 3D structures of Phosphatidylinositol-3-phosphate and Phosphatidylinositol-5-phosphate were drawn using MarvinSketch tool (v23.5, ChemAxon) and exported into SDF formats.

Ligands were subjected to ligand preparation using the LigPrep Wizard in Maestro platform of Schrodinger software. OPLS_2005 force field was applied in order to computationally generate all possible stereoisomers of maximum number 32. Following the creation of stereoisomers, we considered all conceivable conformations of the ligands for the subsequent stages of docking and in-depth analysis [58].

### Molecular Docking

Receptor-grid generation was carried out on the basis of selecting the active site residues of the proteins. Glide program in Schrodinger was used for docking the target protein molecules with all the ligands considering the “Extra Precision (XP)” mode of docking. For getting the best result, number of possible conformation of the protein-ligand complex was taken as one. Ligand interaction module is used for generating the 2D interaction of proteins and ligands. [59]. Docking scores (kJ/mol) for each of the ligands were calculated and the H-bond interaction between ligands and proteins were analysed using LigPlot tool [60].

### Interaction study of PfRKIP WT and mutant proteins with PtdIns(3)P through ITC

ITC measurement were performed to obtain thermodynamic parameter associated with the binding of protein with phospholipid in phosphate buffer on a MicroCal iTC200 [61, 62]. The concentration of the proteins and lipid used for the experiment were 20 µM and 600 µM, respectively. A total volume of 40 µl was titrated from the injection syringe to a sample cell containing 280 µl of the sample at 25 °C. There were a total of 20 injections with each injection of 2 µl of sample titrated into the cell containing buffer in reference cell and each injection was separated by 150 sec intervals to allow the signal to return to baseline. In control experiment, the sample from the syringe was titrated into an ITC cell containing a working buffer. The nonlinear data was fitted with the help of sequential binding model using the MicroCal ORIGIN 7 software supplied by the manufacturer, yielding binding constant (K_avg_), enthalpy change (ΔH) and entropy change (ΔS). ΔG was calculated using relationship, ΔG = ΔH – TΔS.

### Regulation of kinase activity of PfCDPK1 in presence of PfRKIP and its mutant

Regulation of kinase activity of PfCDPK1 by recombinant PfRKIP WT was evaluated using kinase assay as described earlier [27]. Briefly, 100 ng of recombinant PfCDPK1 protein was taken in a kinase assay buffer (50 mM Tris, 50 mM MgCl_2_, 1× phosphatase inhibitor cocktail (Roche Life Science, Indianapolis, IN), 1 mM DTT, 28.3 mM NaCl, 10 mM Ca^2+^) and incubated at 30 °C for 1 h in presence and absence of equal molar ratio of recombinant PfRKIP and myelin basic protein (MBP). Importantly, the reaction was initiated by addition of 100 μM ATPγS as the source of the phosphate group. The reaction was terminated by incubating the reaction mixture with 5 mM EGTA. p-nitrobenzyl mesylate (PNBM, 3.5 mM) (Abcam, Inc., Cambridge, MA) was added into the reaction mixture and incubated at 20 °C for 2 h to allow alkylation of thiophosphorylated serine or threonine residues. Reaction was stopped by adding 1X SDS gel loading dye. The alkylated thiophosphorylated residues (indicating phosphorylation) were detected through Western Blot using anti-thiophosphorylated antibodies (ab92570, abcam)

To study the effect of PfRKIP on the transphosphorylation activity of PfCDPK1 we used Pro-Q^TM^ diamond stain (ThermoFisher Scientific). PfCDPK1 was incubated with different concentrations PfRKIP (2 μg-16 μg) in presence of MBP (10 μg) in the kinase assay buffer (50 mM Tris, 50 mM MgCl_2_, 1× phosphatase inhibitor cocktail (Roche Life Science, Indianapolis, IN), 1 mM DTT, 28.3 mM NaCl, 10 mM Ca^2+^, 100 μM ATP at 30 °C for 1 h. Reaction was stopped by adding 1X SDS gel loading dye. Reaction mixture was run on SDS PAGE and phosphorylation of the protein was detected by staining the gel with the Pro-Q^TM^ diamond stain by following the protocol described in the kit manual. The stained gel was visualised using the ProQ diamond filter in the G:BOX Syngene Chemic Doc (Syngene).

### ELISA for testing the interaction of PfRKIP and mutants with recombinant PfCDPK1

Interaction between recombinant PfRKIP and its mutants with PfCDPK1 was performed using ELISA. Briefly, 96-well microtiter plates were coated overnight with 2 µg/mL of recombinant PfCDPK1 at 4 °C. Subsequently the wells were blocked with 5 % skimmed milk in PBS. Recombinant PfRKIP or its mutants were added in increasing concentrations (0.03125 µg/ml, 0.156 µg/ml, 0.3125 µg/ml, 0.625 µg/ml, 1.25 µg/ml 2.5 µg/ml, 5 µg/ml, 10 µg/ml, and 20 µg/ml) in the binding buffer (50 mM HEPES pH 8.0, 250 mM potassium acetate, 5 mM magnesium acetate; [63] and plates were incubated for 2 h. Bound PfRKIP and its mutants were detected by incubating each well for 1 h with blocking buffer containing anti-PfRKIP antibody (1:2500). The plate was washed 3 times for 5 min each with 1X PBS having 0.01 % Tween 20. Final washing was given with 1XPBS without Tween 20. The wells were subsequently incubated with the blocking buffer containing horseradish peroxidase (HRP)-conjugated anti-rat antibody (1:35000) for 1 h followed by washing with the washing buffer 3 times for 5 min each. The interaction of the proteins was quantified after adding the substrate phenylenediaminedihydrochloride (Sigma Aldrich) plus H_2_O_2_ by measuring the resulting absorbance at 492 nm in the Multiskan SkyHigh Spectrophotometer (ThermoFisher Sientific). All the experiments (n=3) were done in triplicate and mean ± SEM was calculated.

ELISA was also used to test the effect of locostatin on the interaction of PfCDPK1 and PfRKIP. Locostatin is known to modify mammalian RKIP through alkylation of a histidine residue. For conditions requiring testing the effect of modified PfRKIP on interaction with PfCDPK1, the recombinant PfRKIP was pre-incubated with or without different concentrations of locostatin: 50 μM, 100 μM, 200 μM, 400 μM, and 800 μM at 16 °C for 2 h before adding to PfCDPK1 coated wells. To test the effect of locostatin on the interaction of PfCDPK1 and unmodified PfRKIP, the recombinant PfRKIP was not pre-incubated with locostatin rather locostatin was added concomitantly along with other reagents in the interaction buffer. All the experiments were performed in triplicate and mean ± SEM was calculated. The OD_492_ on the Y-axis representing binding of either modified or unmodified PfRKIP with PfCDPK1 in presence and absence of locostatin was plotted with locostatin concentration on the X-axis.

### Locostatin mediated inhibition of Parasite

Highly synchronised *P*. *falciparum* schizonts (38-42 hpi) were adjusted to a final parasitaemia of 0.5 % at 2 % haematocrit and were incubated with different concentrations of locostatin 10 μM, 15 μM, 20 μM, 25 μM, 30 μM, and 35 μM in a 24 well plate. The newly formed rings after 10 h were then scored by Giemsa staining under a light microscope. The percentage ring parasitemia on the Y-axis was plotted against different concentrations of locostatin (μM) on the X-axis from 3 independent biological experiments in duplicate. Statistical significance was calculated using one-way ANOVA in GraphPad Prism 9.

### *In vitro* biotinylation using recombinant PfRKIPBioID2 protein

To verify the biotinylation activity of biotin ligase 2 (BioID2) we have cloned and purified recombinant PfRKIPBioID2 as described above. The proximity biotinylation assay was set with 1 µM of PFRKIPBioID2 protein in the biotinylation buffer (40 mM Tris-Cl pH 8.0, 3 mM ATP, 5.5 mM MgCl_2_, 20 mM KCl) in presence of 2 µM BSA. 10 µM of biotin was added in the reaction mixture requiring biotinylation condition. No biotin was added in the reaction requiring absence of biotin. To test whether recombinant PfRKIP (1 µM) and BSA (1 µM) possess any biotin ligase activity, we included both the proteins (1 µM) separately in the same buffer composition containing biotin. The reaction was incubated for 12 h at 37 °C and stopped by adding 1X SDS dye. The sample were run on SDS-PAGE gel and transferred on the PVDF membrane for Western blotting. Biotinylation of the recombinant proteins was detected using streptavidin-HRP conjugated antibody (1:1000) and blot was developed as described above.

### Co-immunoprecipitation of PfRKIP using Proximity labelling

To test the interaction of PfRKIP and PfCDPK1, we employed a proximity biotinylation assay as described earlier [64]. Briefly, asynchronous cultures of transgenic parasite expressing PFRKIPBioID2V5 and PfRKIPcmyc were grown into 75 cm2 flasks at 2% parasitemia and 4% hematocrit. D-Biotin was added at a final concentration of 200 μM to both the flask and the parasites were allowed to grow for 48 h at 37 °C with replacement of fresh media (± D-Biotin) after 24 h. Parasites were isolated from the RBCs using 0.01 % saponin in PBS and washed multiple times to remove all the haemoglobin and stored at −80 °C till further use. The frozen parasites were lysed with Radio-ImmunoPrecipitation Assay (RIPA) buffer of the following composition: 150 mM NaCl, 1.0 % NP-40, 0.5 % sodium deoxycholate, 0.1 % SDS, 50 mM Tris (pH 8.0), 1 mM phenylmethylsulfonyl fluoride, and 1X protease inhibitor cocktail (complete Mini, EDTA-free, Roche) for 1 h on ice. The tubes were centrifuged at 13,000 × g for 20 min at 4 °C and the soluble extract was diluted three folds with 50 mM Tris-Cl, pH 7.5. The diluted protein extract was incubated with 50μl of Pierce^TM^ Streptavidin Magnetic Beads overnight under rotating condition at 4 °C. After incubation, the beads were separated using DynaMag-Spin (ThermoFisher Scientific) and washed three times with wash buffer and 2 times with distilled water. The beads were boiled in SDS-PAGE sample buffer and processed for Western blotting using anti-V5 and anti-PfCDPK1 antibody.

### Effect of Phosphorylation on the interaction of PfCDPK1 with PfRKIP and modified PfRKIP

To test the effect of phosphorylation on the interaction of PfCDPK1 with PfRKIP and modified PfRKIP we employed ELISA based assay. Recombinant PfCDPK1 was coated on the 96 well plate as has been described above. 800 μM locostatin was used to pre-treat different concentrations of recombinant PfRKIP (21.87, 43.75, 87.5, 175, 350 and 700 nM) at 16 °C for 2 h to ensure its modification (alkylation). DMSO was used as a vehicle control for conditions requiring no modification of PfRKIP. PfRKIP was incubated with PfCDPK1 in the 96 well plate either in presence or absence of 100 μM ATP for 2 h at RT in incubating buffer as described above. Also, 10 mM CaCl_2_ was added in the reaction buffer for activation of PfCDPK1. Bound PfRKIP was detected through anti-PfRKIP antibody. Horseradish peroxidase (HRP)-conjugated anti-rat antibodies (1:35,000) were added in each well for 1 h followed by addition of the substrate, o-phenylenediamine dihydrochloride (Sigma Aldrich) in citrate buffer containing H_2_O_2_. The 96 well plate was read in Multiskan SkyHigh Spectrophotometer (ThermoFisher Sientific) and the absorbance was measured at 492 nm. The O.D. _492 nm_ values on the Y-axis were plotted against different concentrations of unmodified and modified PfRKIP (nM) on the X-axis. The statistical significance was calculated using paired t-test in GraphPad Prism 9 for 3 independent biological experiments done in duplicates.

## FIGURE LEGENDS

**Figure S1. PfRKIP antibodies detect recombinant protein and PfRKIP in the parasite lysate. a)** Antibodies generated in-house against PfRKIP readily detect the recombinant protein in Western blot. Recombinant PfRKIP was separated on SDS-PAGE and processed for Western blotting with anti-PfRKIP antibodies and preimmune sera. Band corresponding to the recombinant PfRKIP was specifically detected by α PfRKIP Ab whereas preimmune sera did not show any reactivity. **b)** PfRKIP is detected in parasite lysate by α PfRKIP Ab. Parasite lysate prepared from late asexual stage was separated on SDS-PAGE and processed for Western blot with α PfRKIP Ab and preimmune sera. Band corresponding to PfRKIP (~ 21.5 kDa) was detected specifically with the α PfRKIP Ab.

**Figure S2. PfRKIP is localized in mature gametocytes.** Glass slide containing mature gametocytes of *P*. *falciparum* were incubated with anti-PfRKIP antibodies followed by staining with a secondary antibody conjugated with AlexaFluor 488. Stage IV (bottom), stage V male (top) and stage V female (middle) gametocytes stained for PfRKIP (green) are shown. The parasites were counterstained with 4′,6-diamidino-2-phenylindole (DAPI), a nuclear stain. Merge of DAPI and PfRKIP along with Differential Interference Contrast (DIC) are shown. White scale bar is 5 μm in length.

**Figure S3. *In silico* docking of ligands with PfRKIP and HsRKIP.** Three different ligands **i)** phosphorylethanolamine, **ii)** phosphate and **iii)** phosphatidic acid were docked with **a)** PfRKIP (PDB ID: 2R77) modelled at pH 7.5 (PfRKIP PROPKA 7.5), **b)** PfRKIP at pH 5.5 (PDB ID: 2R77) and **c)** *Homo sapiens* RKIP (HsRKIP) at pH 6.5 (PDB ID: 1BD9). The 3D structures of corresponding proteins docked with ligands (orange and green sticks) and LigPlot are shown. The interacting residues of PfRKIP are represented by cyan color in the 3D model.

**Figure S4. PfCDPK1 immobilized on 96 well multiplate is enzymatically active.** PfCDPK1 immobilized on the surface of 96 well multiplate was incubated with myeline basic protein (MBP) in a kinase assay buffer either in presence or absence of locostatin and ATP. Only in presence of ATP, the phosphorylation of MBP was detected. Enzymatic activity of PfCDPK1 was also evident in the presence of locostatin in the kinase assay buffer.

**Figure S5. Locostatin modified recombinant PfRKIP show less binding with PfCDPK1 in presence of ATP.** Recombinant PfRKIP pre-incubated with locostatin was allowed to interact with recombinant PfCDPK1 either in presence or absence of ATP. In presence of ATP, the modified PfRKIP show less binding with PfCDPK1. Concentration of recombinant PfRKIP (in nM) on the X-axis is plotted against O.D. _492 nm_, representing binding of PfRKIP with PfCDPK1 in presence and absence of ATP, on the Y-axis from three independent biological experiments performed in duplicate. Error bars represent standard error of mean. Statistical significance was calculated using GraphPad Prism 9. P value= 0.0040, paired t-test.

**Figure S6. Pre-incubation of PfRKIP with locostatin is essential for increased binding of modified PfRKIP with PfCDPK1.** Inclusion of recombinant PfRKIP along with locostatin (without pre-incubation) in the kinase assay reaction with PfCDPK1 does not show any change in binding of PfRKIP with PfCDPK1 compared to without locostatin condition. PfRKIP modified by pre-incubation with locostatin is taken as a positive control. The graph is plotted with concentration of locostatin (in μM) on the X-axis against O.D. _492 nm_, representing binding of PfRKIP with PfCDPK1 in presence and absence of locostatin, on the Y-axis. The graph is plotted from data obtained from 2 independent biological experiments performed in duplicate. GraphPad Prism 9 is used for plotting the histogram.

**Figure S7. Verification of enzymatic activity of biotin ligase 2 and presence of episomal plasmid carrying *pfrkip* tagged with biotin ligase 2-V5 in the transfected parasites.** a) Chimeric recombinant PfRKIP tagged at the C-terminus with biotin ligase 2 was expressed and purified from *E*. *coli*. The chimeric protein was used to biotinylate bovine serum albumin (BSA) in an *in vitro* assay in presence and absence of biotin. Recombinant PfRKIP was used as a negative control. Left panel shows the Western blot probed with streptavidin-HRP conjugated protein. Right panel represents the reactions mixtures separated on SDS-PAGE and stained with Coomassie Blue R250 dye. M-unstained protein standard. **b)** A diagnostic PCR was used to test the presence of episomal plasmid containing biotin ligase 2-V5 tagged *pfrkip* gene. Primer pair pDC2A/ pDC2B specific to the plasmid sequence flanking the site of insertion of *pfrkip* was used to amplify target sequence from *pfrkip-bioid2v5* and *pfrkip-cmyc* parasites. Amplicons of desired sizes (1525 bp and 832 bp) were obtained in the *pfrkip-bioid2v5* and *pfrkip-cmyc* parasites, respectively. WT parasites were used as a negative control. M-GeneRuler DNA Ladder Mix.

## ACKNOWLEDGEMENTS

We thankfully acknowledge Josh Beck for pm2gt-hsp101-bioid2 plasmid, Jacobus Pharmaceutical Company, Inc & Kristin D. Lane for WR99210, Judith L. Green & Anthony A. Holder for the anti-PfCDPK1 antibodies, and Pankaj Kesari & Pramod C. Rath for HEK293T cDNA. Financial support from the Department of Biotechnology (#BT/PR28256/MED/29/1313/2018 and BT/PR38411/GET/119/311/2020) to AB is also gratefully acknowledged. Financial support from PAC-JNU-DST-PURSE-462/Department of Science and Technology, Ministry of Science and Technology is also acknowledged. The animal facility of JNU, Central Laboratory Animal Resources (CLAR) is acknowledged for generation of antisera against recombinant PfRKIP protein in rats. The funding agency has no role in the preparation and decision to publish the work. MS is a recipient of JRF-SRF Fellowship from the Council for Scientific and Industrial Research. We also acknowledge all the lab members who assisted and provided valuable feedback during the course of the study. Inability to include work from other colleagues on the subject is regretted.

