## supporting figures for "*Plasmodium falciparum* Raf kinase inhibitor is a lipid binding protein that interacts with CDPK1 and regulates its activity in asexual blood stage"

### Figure S1

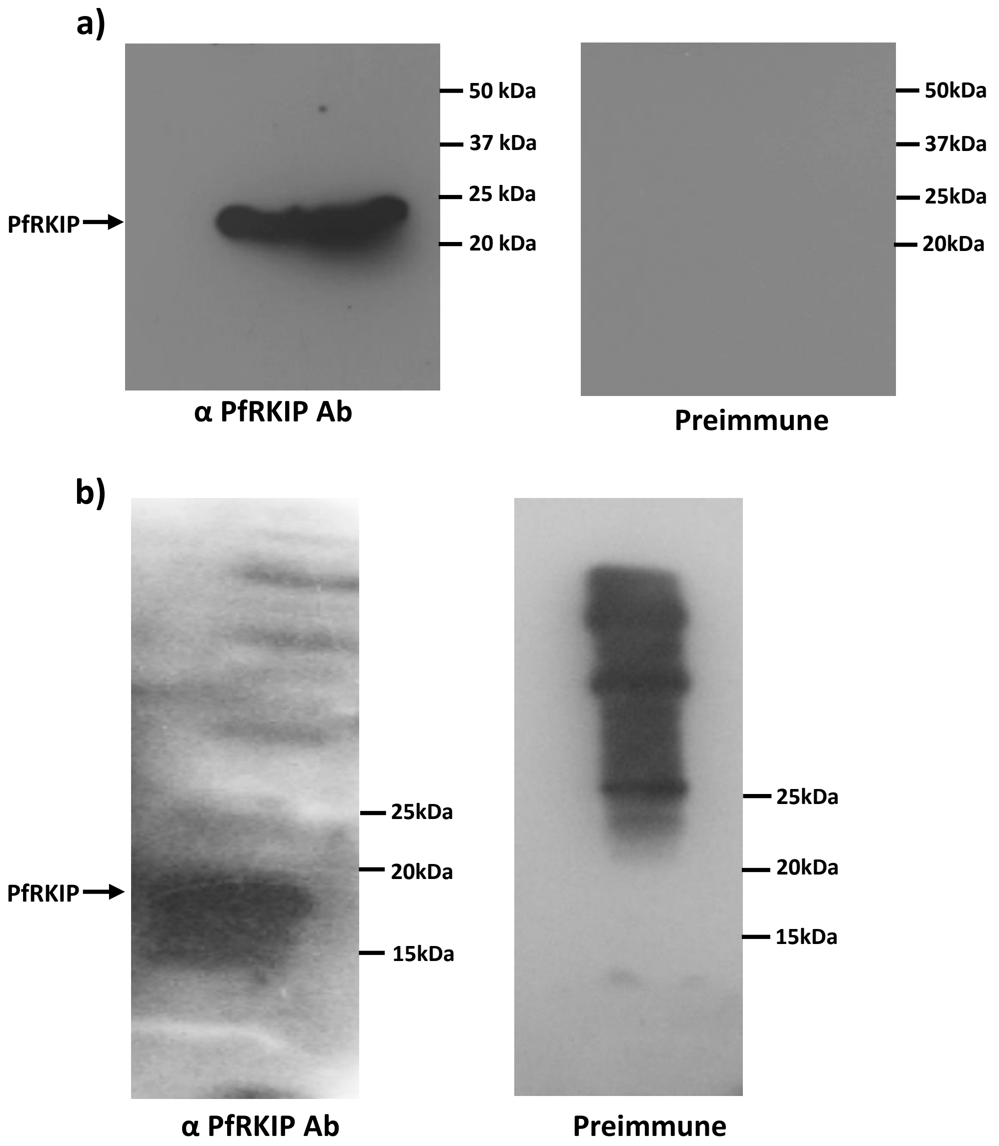

a)
PfRKIP
50 kDa
37 kDa
25 kDa
20 kDa
α PfRKIP Ab
50kDa
37kDa
25kDa
20kDa
Preimmune
b)
PfRKIP
25kDa
20kDa
15kDa
α PfRKIP Ab
25kDa
20kDa
15kDa
Preimmune

Figure S2

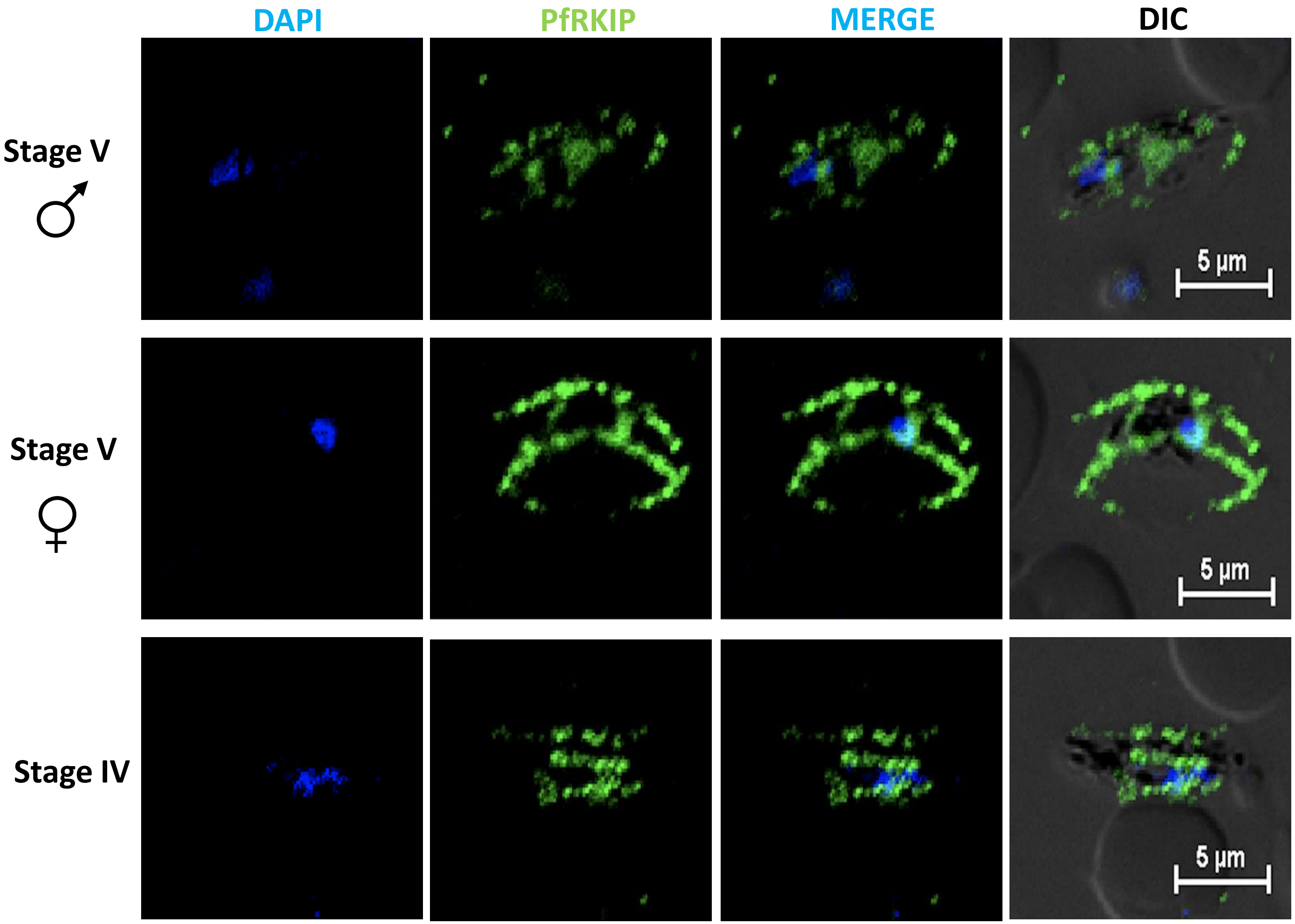

Figure S3

**a)**

**i)**

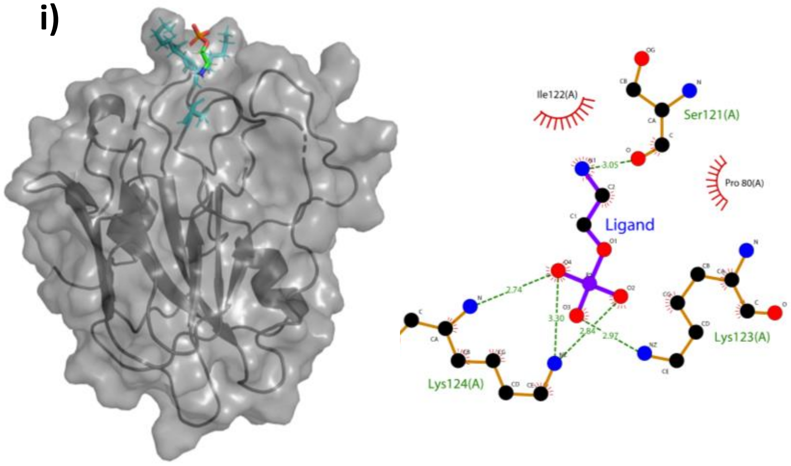


**PfRKIP PROPKA 7.5 + Phosphoryl_ethanolamine**

**ii)**

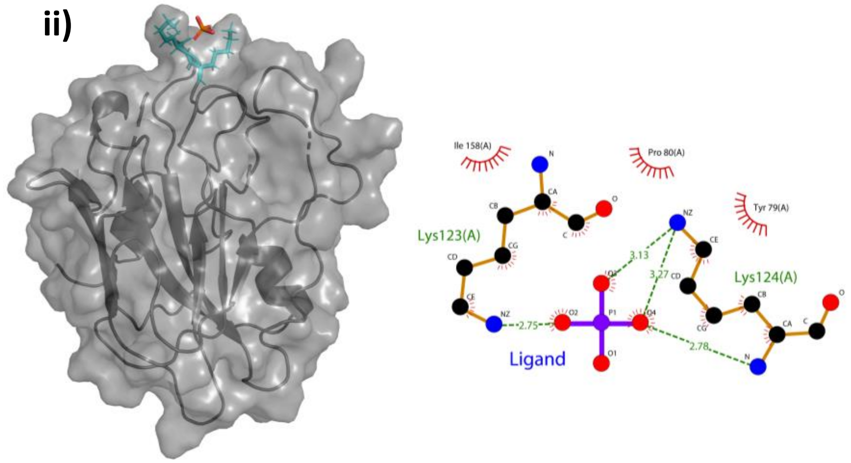


**PfRKIP PROPKA 7.5 + Phosphate**

**iii)**

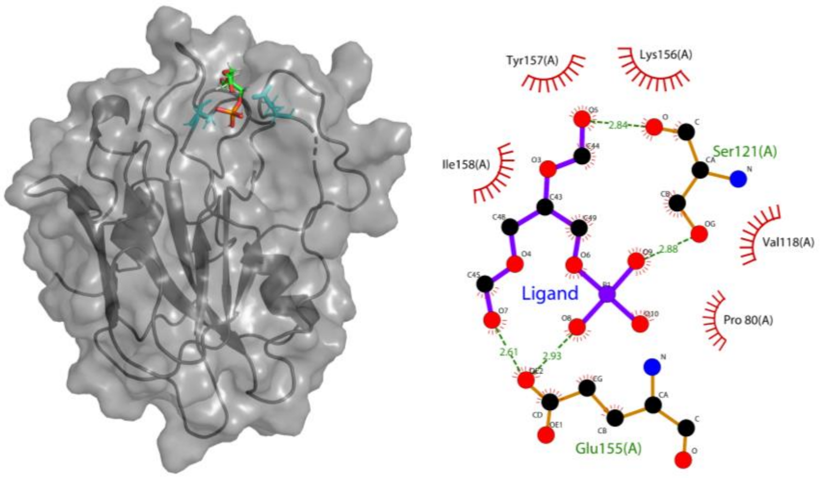


**PfRKIP PROPKA 7.5 + Phophatidic_acid**

Figure S3

**b)**

**i)**

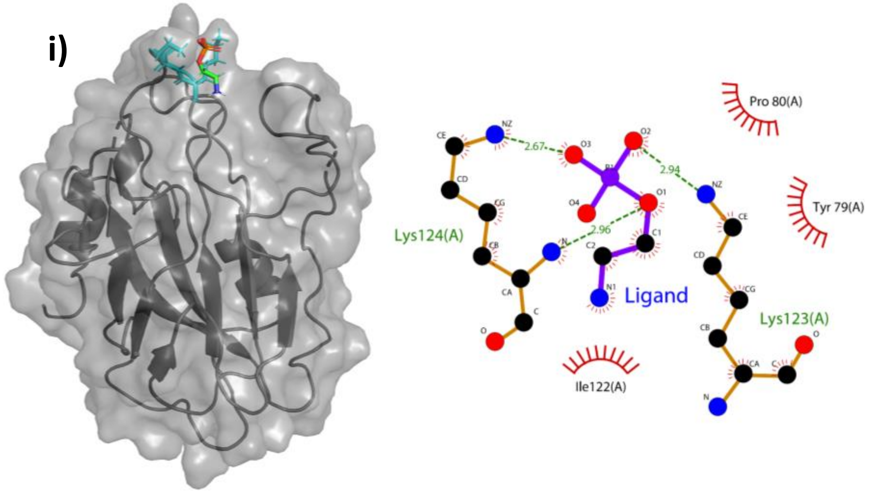


**PfRKIP (pH 5.5) + Phosphoryl_ethanolamine**

**ii)**

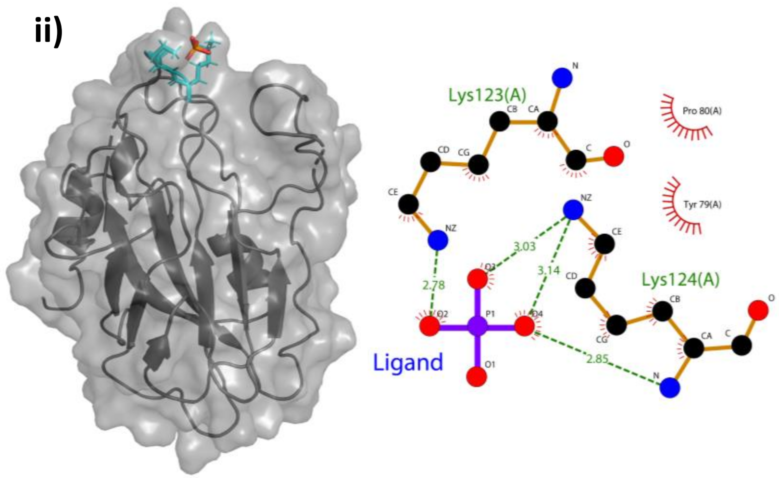


**PfRKIP (pH 5.5) + Phosphate**

**iii)**

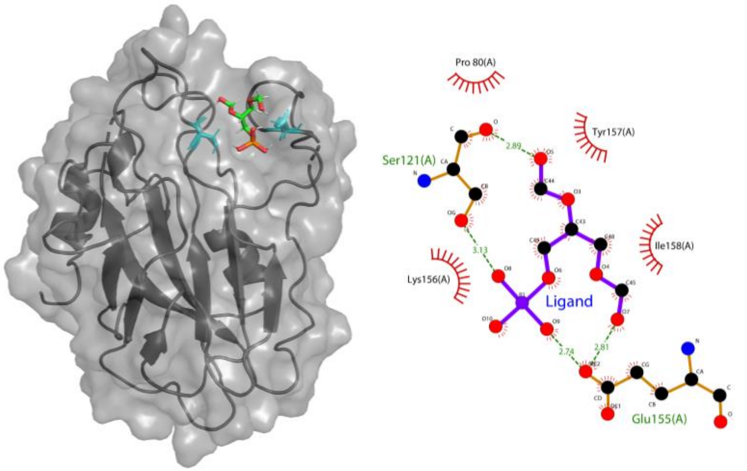


**PfRKIP (pH 5.5) + Phophatidic_acid**

Figure S3

c)

i)

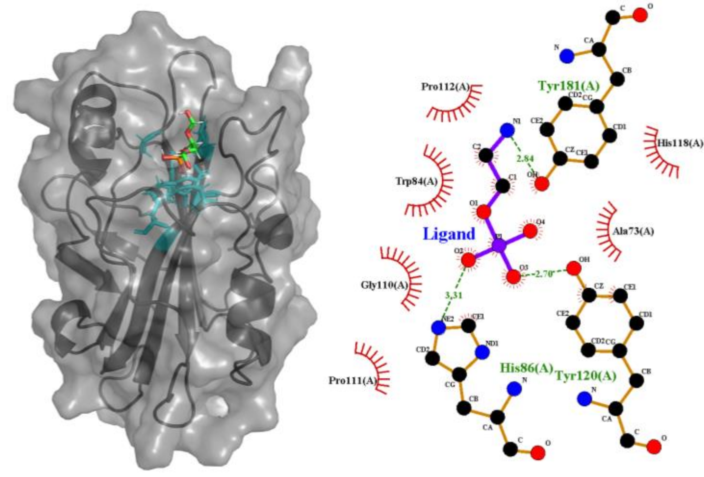


HsRKIP (pH 6.5) + Phosphoryl_ethanolamine

ii)

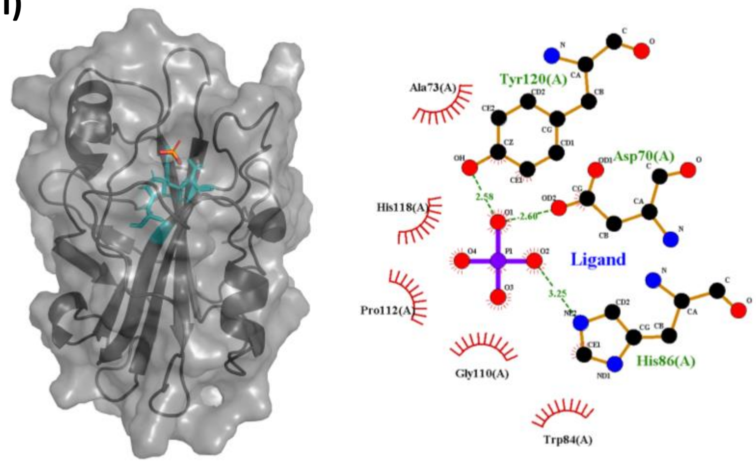


HsRKIP (pH 6.5) + Phosphate

iii)

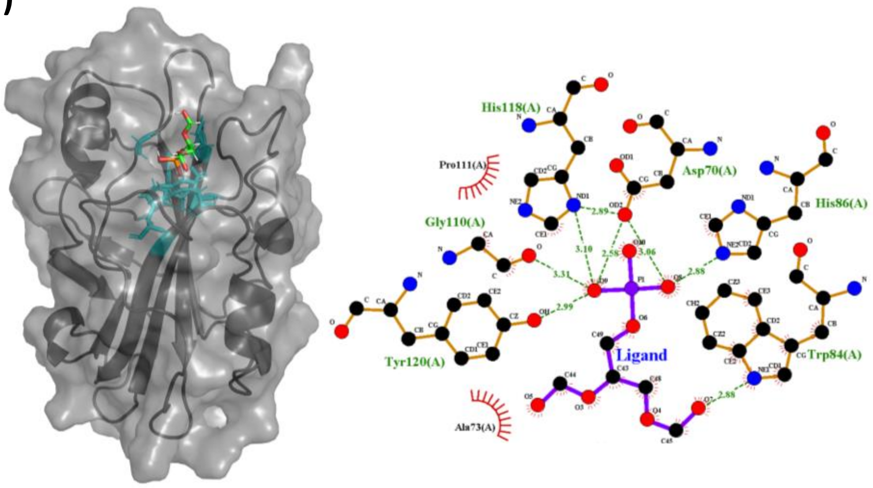


HsRKIP (pH 6.5) + Phophatidic_acid

Figure S4

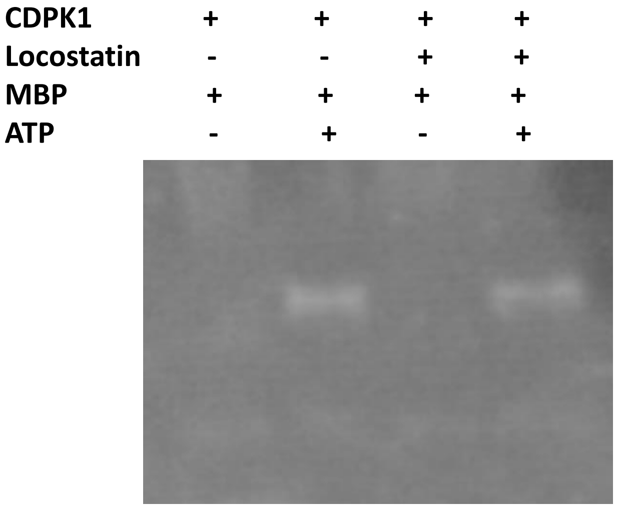

Figure S5

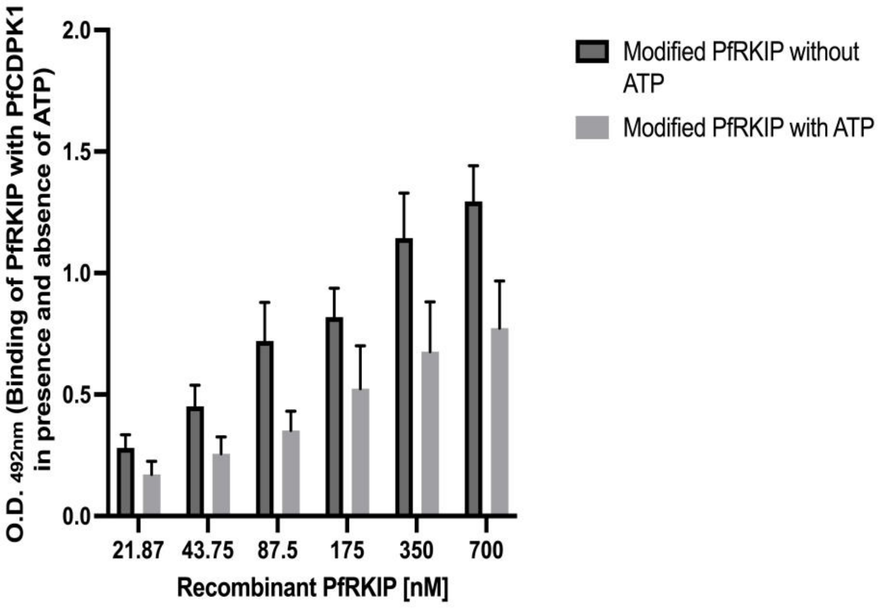

O.D. 492nm (Binding of PfRKIP with PfCDPK1 in presence and absence of ATP)
2.0
1.5
1.0
0.5
0.0
21.87
43.75
87.5
175
350
700
Recombinant PfRKIP [nM]
Modified PfRKIP without ATP
Modified PfRKIP with ATP

Figure S6

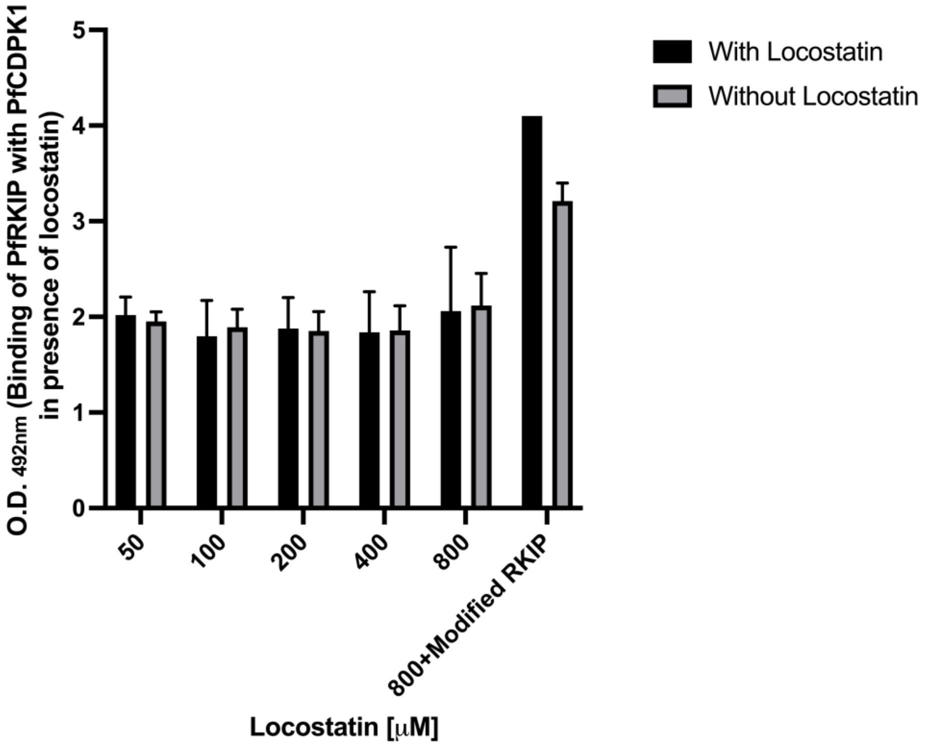

O.D. 492nm (Binding of PfRKIP with PfCDPK1 in presence of locostatin)
5
4
3
2
1
0
50
100
200
400
800
800+Modified RKIP
Locostatin [μM]
With Locostatin
Without Locostatin

Figure S7

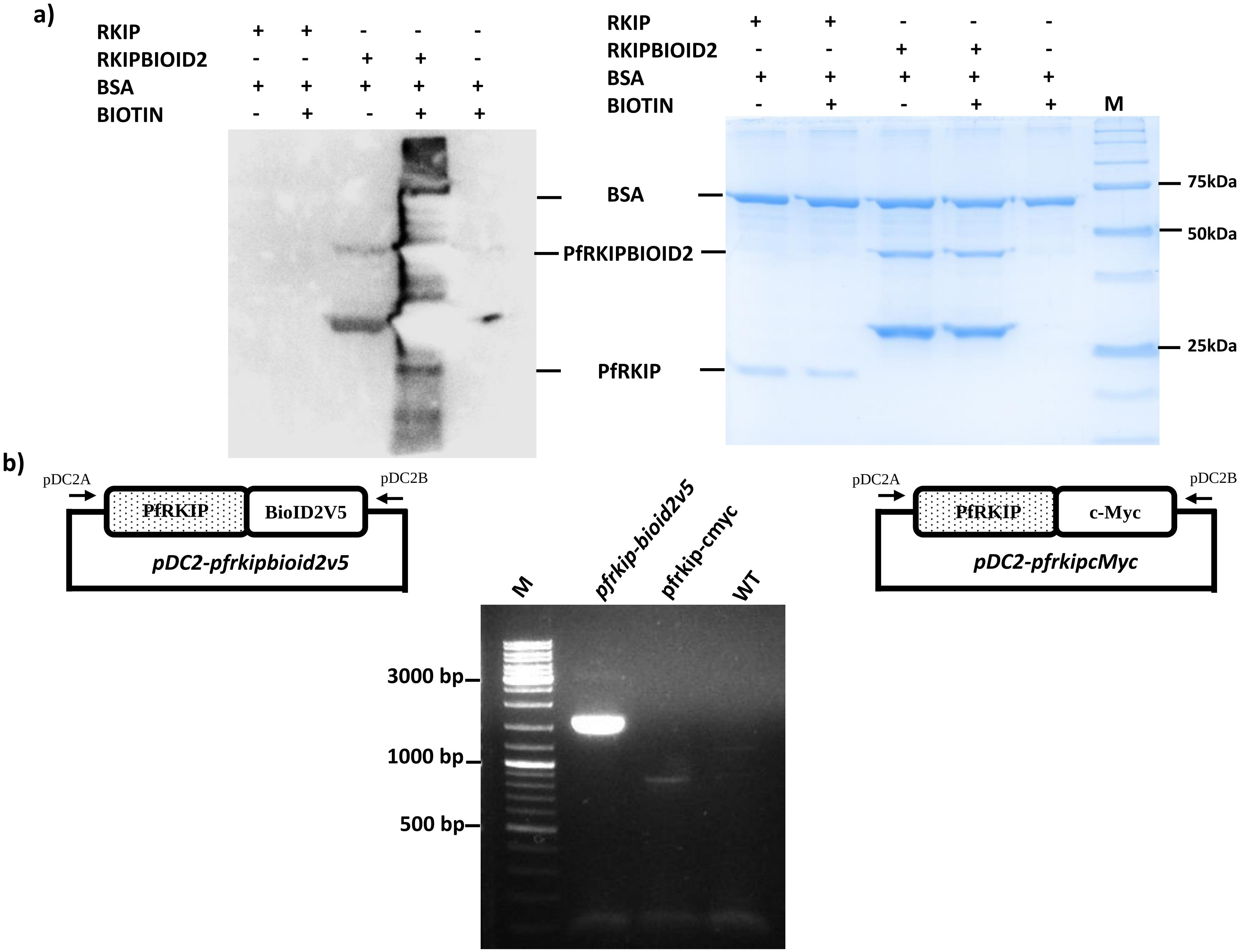
a)
RKIP + + - - -
RKIPBIOID2 - - + + -
BSA + + + + +
BIOTIN - + - + +
BSA
PfRKIPBIOID2
PfRKIP
RKIP + + - - -
RKIPBIOID2 - - + + -
BSA + + + + +
BIOTIN - + - + +
M
75kDa
50kDa
25kDa
b)
pDC2A
pDC2B
PfRKIP
BioID2V5
pDC2-pfrkipbioid2v5
M
pfrkip-bioid2v5
pfrkip-cmyc
WT
3000 bp
1000 bp
500 bp
pDC2A
pDC2B
PfRKIP
c-Myc
pDC2-pfrkipcMyc
